# Tumors Exploit Dedicated Intracellular Vesicles to Program T cell Responses

**DOI:** 10.1101/691873

**Authors:** Edward W. Roberts, Megan K. Ruhland, En Cai, Adriana M. Mujal, Kyle Marchuk, Casey Beppler, David Nam, Nina K. Serwas, Mikhail Binnewies, Matthew F. Krummel

**Affiliations:** Department of Pathology, University of California San Francisco, San Francisco, CA 94143, USA; Beatson Institute for Cancer Research, Glasgow, UK; Immunology Program, Memorial Sloan Kettering Cancer Center, New York, NY, 10065, USA; Biological Imaging Development CoLab, University of California, San Francisco, CA 94143, USA; Pionyr Immunotherapeutics, San Francisco, CA 94107, USA

## Abstract

In order to drive productive tumor-infiltrating lymphocyte (TIL) function, myeloid populations must direct antigens to the lymph node, including to resident antigen-presenting cells (APCs) that have never touched the tumor. It has long been supposed that APCs trade antigens with one another, but the dominant cell biology underlying that remains unknown. We used *in vitro* and *in vivo* assays together with lattice light sheet and multiphoton imaging to show that myeloid cells carry tumor antigen-laden vesicles that they ‘trade’ with one another as they reach distant sites. This accounts for the majority of antigen displayed to T cells and provides tumors with a mechanism to access APCs that differentially direct T cell activation away from memory phenotypes. This work defines efficient cell biology that drives the first steps of TIL generation and represents a new frontier for engineering tumoral immunity.

## Main Text

Immune responses to cancer require the movement of tumor-derived antigens into the tumor draining lymph node (tdLN) where T cell priming takes place on a diverse set of dendritic cell (DC) populations. Previous work has shown that migratory DC populations are required to carry tumor antigens to the tdLN and more broadly to disseminate tumor antigens to tdLN resident DC populations, but the biological mechanism for this has been unclear(*1*). Peptides injected in the context of vaccination can directly drain to the LN for sampling by resident myeloid cells(*2, 3*) whereas migratory cells are required to carry antigen to the LN in other physiological challenges such as infection or tolerance, and the antigens are then subsequently acquired by resident DC(*4*– *10*). In the context of an actively growing tumor, much less is known; competing hypotheses from other systems suggest migratory DC may: 1. Undergo apoptosis and be taken up by resident cells(*7*); 2. Secrete exosomes containing antigen for subsequent stochastic re-uptake(*11*); 3. Secrete soluble antigens into the LN interstitium(*12*) or 4. Transfer entire peptide-MHC complexes, in a process termed cross-dressing and likely most important for memory T cell activation(*13*).

To track dissemination of tumor proteins, B16F10 cells were modified to express the pH-stable fluorophore ZsGreen (B16ZsGreen), a fluorophore that persists much longer than others in intracellular compartments and allows long-term optical tracking of proteins(*1*). These tumors were then grown in mice expressing membrane bound tdTomato (mT) on all host cells (**Fig. 1A**) and both the tumor and the draining lymph node (tdLN) were examined. This revealed that host cells in the tumor microenvironment (TME) contained significant quantities of tumor-derived ZsGreen, and this material was consistently observed as vesicular puncta (**Fig. 1A**) and never as uniform cytoplasmic distributions. By flow cytometry, digested B16ZsGreen tumors showed significant ZsGreen uptake by host CD45^+^ cells (**Fig. 1B**) and uptake was penetrant: 100% of tumor associated macrophages (TAMs) and conventional dendritic cell (DC) type 1 (cDC1) and type 2 (cDC2) and ∼60% of monocytes contained significant amounts of ZsGreen (**Fig. 1C**, see **Fig. S1** for gating strategy). To further analyze these vesicles and to determine whether this ZsGreen^+^ compartment accurately represented the localization of germline-encoded tumor antigens in these cells, ZsGreen^+^ myeloid cells were sorted from the TME, stained for melanoma tumor-associated antigens, gp100 and tyrosinase, and imaged by confocal microscopy (**Fig. 1D**). These known tumor antigens were also packaged as discreet puncta, consistent with vesicles. Importantly, within the TME, >75% of ZsGreen containing vesicles contained at least one of these tumor antigens (**Fig. 1D**). For the remainder, we noted that gp100 and tyrosinase are typically located within vesicles in the tumor cells and therefore may be variably distributed after sampling, as compared to cytosolic ZsGreen. We conclude that the host compartment containing ZsGreen also faithfully reports other tumor antigens.

**Fig. 1.**
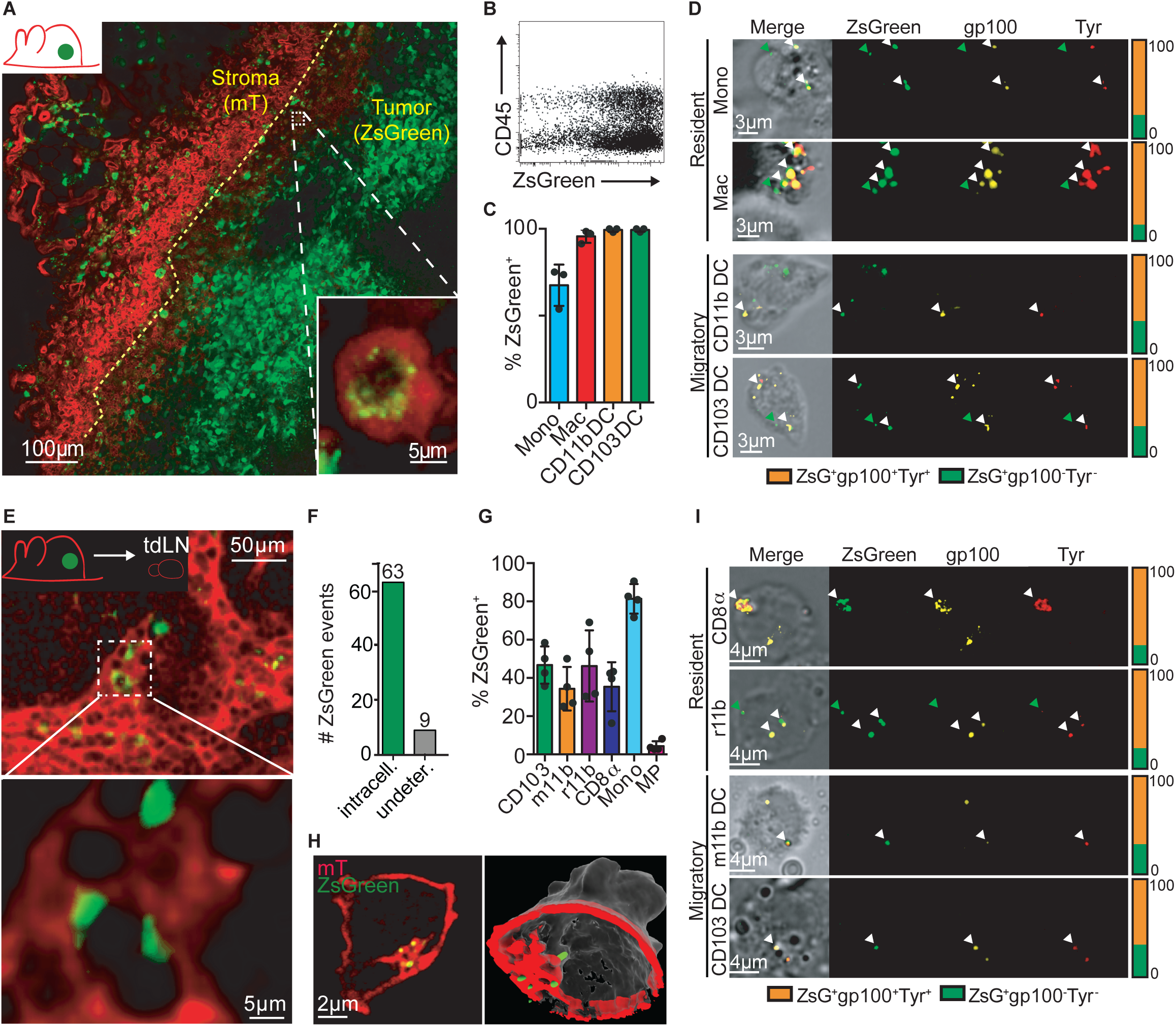
Visualization of tumor antigens within discrete migratory and resident myeloid compartments. **(A)** B16zsGreen tumors injected into membrane-tdTomato (mT) expressing mice were isolated and imaged by confocal micrcroscopy to detect ZsGreen^+^ puncta within host mT ^+^cells of the tumor microenvironment. Representative image. Scale bar = 100µm. Inlay image is higher magnification of intracellular ZsGreen^+^ puncta within mT ^+^host cells. Representative image. Scale bar = 5µm. (**B**) Representative flow cytometry plot showing composition of CD45^+^, ZsGreen^+^ cells within the tumor microenvironment of B16ZsGreen tumors. (**C**) Quantification of tumor-derived ZsGreen within myeloid cells of the tumor. Plot shows mean frequency +/−SEM. Data are representative of 3 independent experiments. (**D**) ZsGreen^+^ myeloid cells from B16ZsGreen tumors stained for tumor-associated antigens, tyrosinase (Tyr, red) and gp100 (yellow). Frequency of colocalization for tumor-derived ZsGreen with tyrosinase and gp100 is quantified (right) for each cell type. Representative images. Arrows indicate examples of colocalized puncta. Scale bars = 3µm. (**E**) Tumor draining lymph nodes (tdLN) were isolated, fixed and cleared prior to confocal imaging to detect ZsGreen puncta within host mT cells of the tdLN. Representative image. White box inset (top) shows position of zoomed+image (bottom). Top image scale bar = 50µm. Bottom image scale bar = 5µm. (**F**) Quantification of ZsGreen localization pattern observed in tdLN (see image in Fig.S2). Plot shows number of ZsGreen events counted that were contained within an intact tdTomato^+^ cell membrane (intracellular) compared to those that were not (undetermined). (**G**) Flow cytometric quantification of tumor-derived ZsGreen within myeloid cells and microparticles (MP) of the tdLN. migratory CD11b (m11b), resident CD11b (r11b). Plot shows mean frequency +/−SEM. Data are representative of 3 independent experiments. (**H**) Lattice light sheet imaging of sorted dendritic cell (DC) from B16ZsGreen tdLN. Left image shows z-slice of cell with tumor-derived ZsGreen evident as puncta. Representative image. Scale bar = 2µm. Right is same cell with surface rendering to show single z-slice of membrane (red) with remaining depth of z plane in grey. Total 3D volume of ZsGreen is surfaced (green) to illustrate host membrane encapsulation of intracellular puncta. Image rendered using ChimeraX software. (**I**) ZsGreen^+^ myeloid cells were sorted from B16ZsGreen tdLN and stained for tumor-associated antigens, tyrosinase (Tyr, red) and gp100 (yellow). Frequency of colocalization for tumor-derived ZsGreen with tyrosinase and gp100 is quantified (right) for each cell type. Arrows indicated examples of colocalized puncta. Representative images. Scale bar = 4µm.

When we used conventional confocal imaging to examine mT^+^ host tdLNs, ZsGreen was again observed within cells as puncta (**Fig. 1E**). We rarely observed ZsGreen material that was not within these intracellular vesicles (**Fig. 1F, S2**) and no accumulation was observed in the subcapsular sinus where microparticles would be expected to accumulate (**Fig. S2**). The frequency of cells that were ZsGreen^+^ in the tdLN was only about 40% of both migratory and resident cDC1 (migratory CD103^+^ DC and resident CD8α^+^ DC) and cDC2 (migratory CD11b^+^ DC and resident CD11b^+^ DC), as compared to nearly 100% in the tumor (**Fig. 1C, G**). Particularly in mice bearing very large tumors, we often also found plentiful ZsGreen^+^ monocytes but again only identified relatively small numbers of cell-free ZsGreen^+^ subcellular sized microparticles (defined as lacking nuclei and weakly scattering by flow cytometry) (**Fig. 1G**). We sorted cells from the tdLN and analyzed them live, by high-resolution lattice light sheet microscopy (LLS) in order to resolve the puncta and minimize background fluorescence. All of the detectable ZsGreen signal localized to vesicular structures (ZsGreen signal surrounded by mT^+^ membrane shown in **Fig. 1H**, image rendering (right) highlights ZsGreen puncta membrane encapsulation by showing 3D depth in z (cell volume, grey) see also **Movie S1**). As in cells derived from the TME, the majority of ZsGreen^+^ puncta, including those in resident DC populations, also contained the other canonical tumor-associated antigens, again suggesting a common vesicular compartment for internalized antigen (**Fig. 1I**).

In previous studies using genetically deficient mice and also confirmed with mixed bone marrow chimera experiments, CCR7 was shown to be required for DC migration from the tumor to the tdLN and for the subsequent ability to activate T cells(*1, 14*). We extended that finding here, showing that ZsGreen^+^ cells and microparticles were virtually absent in the tdLN of *Ccr7*^*-/-*^ mice (**Fig. S3A**). In contrast, in mice lacking CCR2, a chemokine required for monocyte migration, ZsGreen levels were comparable to controls (**Fig. S3B**). In wildtype animals, detailed study of ZsGreen levels in migratory DC of the tdLN demonstrated a 10-fold reduction in fluorescence intensity as compared to tumor-resident counterparts (**Fig. 2A**). Likewise, ZsGreen^+^ vesicle number per cell in the migratory DC decreased by over half when looking at migratory DC in the tdLN compared to the tumor (**Fig. S3C**). Furthermore, ZsGreen^+^ resident DC were more numerous and of comparable brightness to migratory DC in the tdLN (**Fig. 2B**). These observations are consistent with a fixed amount of fluorescence being actively redistributed, as vesicular packets, into a larger number of recipient cells in the tdLN. Thus, we considered the possibility of directed handoff to resident cells, leading to the dilution of the fluorescence in migratory populations. Such a mechanism would represent the cellular equivalent of the ancient mancala ‘sowing’ games in which rules determine how and when ‘seeds’ are moved from one players compartments into the compartments of another.

**Fig. 2.**
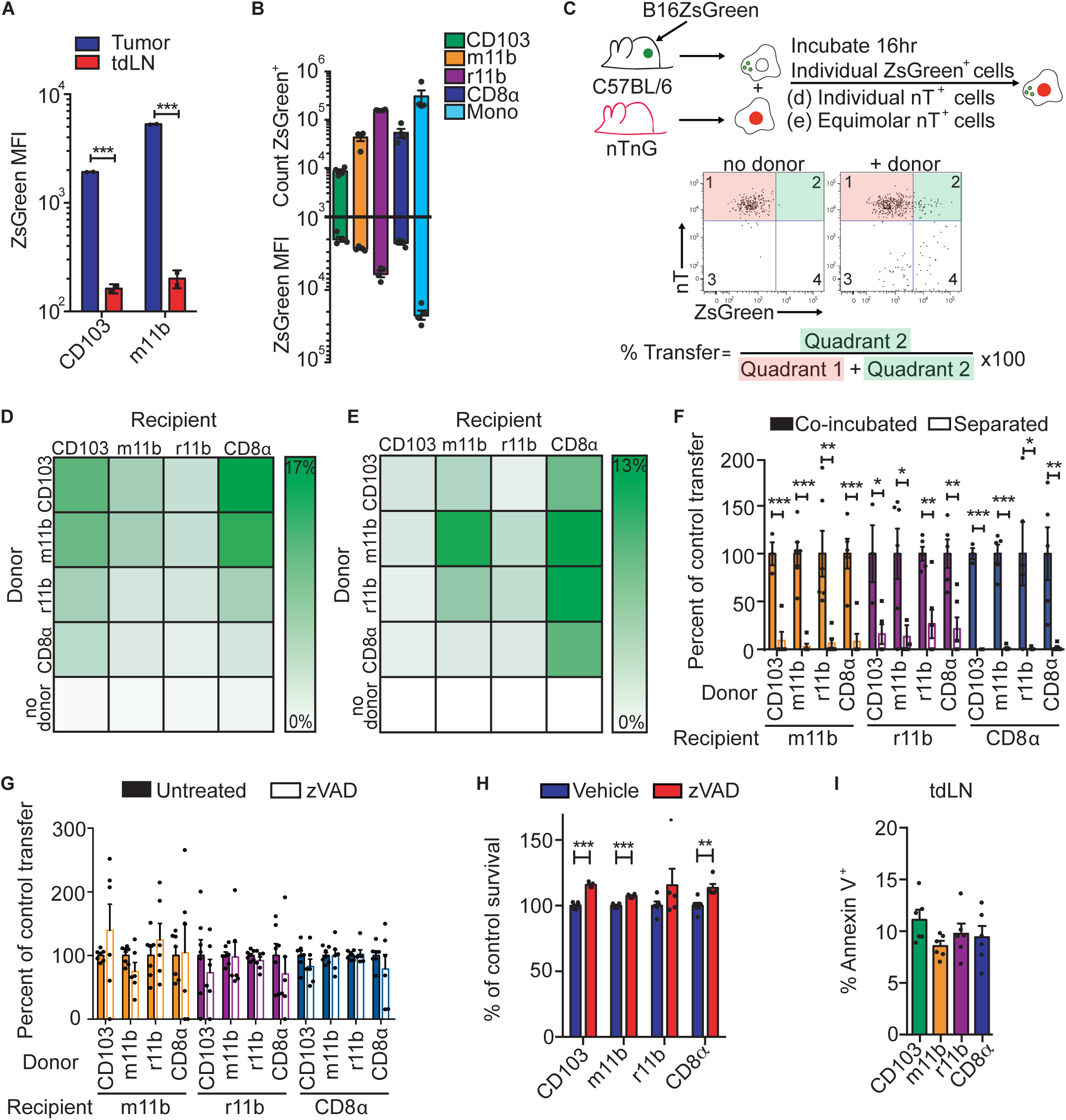
Pathway of tumor antigen transit and transfer. **(A)** Quantification of the mean fluorescence intensity (MFI) of ZsGreen within migratory dendritic cell (DC) types in B16ZsGreen tumors and matched tumor draining lymph nodes (tdLN). Plot shows MFI +/−SEM. Representative of 2 independent experiments. (**B**) Quantification of the number of ZsGreen^+^ cells and the MFI of ZsGreen within myeloid cells of the tdLN. migratory CD11b (m11b), resident CD11b (r11b). Plots show count and MFI +/−SEM. Representative of 3 independent experiments. (**C**) *In vitro* antigen transfer assay. Donor ZsGreen ^+^DC are isolated from B16ZsGreen tdLN and recipient DC are isolated from LN of nuclear-tdTomato (nT) mice. Donor and recipient DC are incubated together as in D-E. Flow plot shows method for calculating % of recipient cells acquiring antigen. (**D-E**) ZsGreen^+^ donor cells were isolated from tdLN of B16ZsGreen mice and (**D**) co-cultured with individual recipient DC at ratio 1:1 or (**E**) co-cultured with an equal ratio mix of all four recipient DC. Heat maps show mean frequency of ZsGreen^+^ nT ^+^recipients after co-culture +/−SEM. Representative of 4-5 independent experiments. (**F**) Setup as in E with the addition of 3µm pore transwell. Error bars represent +/−SEM. Representative of 3 independent experiments. (**G**) Setup as in e conducted in the presence of zVAD. Representative of 3 independent experiments. (**H**) Survival of zVAD-treated DC relative to vehicle-treated DC *in vitro*. Plot shows mean relative % survival of zVAD-treated cells compared to vehicle. Error bars represent +/−SEM. Representative of 3 independent experiments. (**I**) Annexin V staining for DC types in the tdLN. Plot shows mean % of Annexin V^+^ apoptotic cells. Error bars represent +/−SEM. Representative of 3 independent experiments. In all graphs *p < 0.05, **p < 0.01,***p < 0.001.

Consequently, we developed an *in vitro* ‘mancala assay’ wherein *in vivo*-loaded ZsGreen^+^ ‘donor’ DC from tdLN were incubated with nuclear tdTomato (nT)^+^ ‘recipient’ DC. After 16 hours, we used flow cytometry to determine whether, and under what conditions, tumor-derived material transferred from one cell type to another (**Fig. 2C**). Using a single type of recipient cell, we observed a pattern wherein either population of migratory DC served as an equally proficient donor and cDC1—especially resident CD8α^+^ DC—were consistently the best recipients (**Fig. 2D**). In our assays, we found that all populations formed contacts at similar levels with donors, as assessed by measuring doublets, suggesting that it was the actual transfer and not a contact-event that was regulated (**Fig. S4A-B**). We also performed a competitive version of the mancala assay, in which all four recipient cell types were present in the same well, to assess dominance. Here, there was less consistency in terms of which donor cell type produced the highest degree of transfer, but the resident cDC1 CD8α^+^ DC always ultimately accumulated the majority of the material, thus serving as the best recipient, by the conclusion of the assay (**Fig. 2E**). While both the individual and the competitive mancala assays ordained the CD8α^+^ DC the superior recipient *in vitro*, this may be tempered *in vivo* where the localization and number of particular DC subsets likely play a role and we indeed see resident CD11b^+^ DC as active participants in tumor antigen uptake *in vivo* (**Fig. 2B**). Beyond the cellular level, we next used this assay to more completely address the vesicular nature of the ‘passed’ antigen. To do this, we performed the mancala assay using ZsGreen-loaded DC sorted from the LN of mT^+^ mice bearing B16ZsGreen tumors. The ZsGreen^+^,mT^+^ DCs were co-cultured with unlabeled, naïve, congenically marked, recipient DC (**Fig. S5A**). After 16hr, the CD45.1^+^ZsGreen^+^ recipient DCs were sorted, cell surface-stained with anti-CD45 and imaged by LLS. The high-resolution imaging revealed intracellular ZsGreen^+^ puncta enveloped by donor DC mT^+^ membrane demonstrating that passed antigen is handed off to the recipient DC in packets, likely the vesicles themselves (**Fig. S5B**).

Next, we varied the mancala assay to provide insight into mechanism. First, we found that the entire process of transfer was abrogated when populations were separated by a transwell (**Fig. 2F**) suggesting, but not proving, that antigen is not passed in soluble form nor in exosomes in this system, in agreement with previously published data using *in vitro* generated bone marrow derived dendritic cells (BMDC)(*15*). We next applied a pharmacological perturbation. Here, we eliminated migratory cDC1 from the recipient pool as they were previously determined to be poor recipients and furthermore were more difficult to purify in sufficient numbers for this assay. Amongst many inhibitors tested, we found that PI3K class I (GDC0941) or III (VPS34-IN1) inhibition, which prevents phagocytosis, strongly inhibited CD8α^+^ cDC1 uptake, and produced less pronounced effects on the other recipient types (**Fig. S6A-B**). CD8α^+^ DC have been previously described as being more phagocytic than cDC2, perhaps also explaining their dominance in antigen uptake *in vitro* (**Fig 2E)**(*9, 16, 17*). Overall, this suggests that different DC may rely on different mechanisms for antigen uptake.

We also applied this assay to specifically interrogate the role of DC apoptosis in antigen transfer. We first added zVAD, which blocks caspase activity, and saw no effect on transfer for any combination (**Fig. 2G**) despite increases in survival of donor DC subsets (**Fig. 2H**). However, unlike the BMDC used in previous studies, DC from tdLN showed low rates of apoptosis overall suggesting that high rates of DC death both *in vitro and in vivo* may not represent a prominent source of antigen transfer for *in vivo* DC populations (**Fig. 2I**).

Given that transwells blocked antigen transfer, we explored the hypothesis that vesicle transfer required direct cell-cell contact. To date, myeloid-myeloid interactions have not been well documented and so we devised a real-time, 2-photon, imaging strategy to study these in the interfollicular T cell zone of the tdLN of mice bearing B16ZsGreen tumors. To do this in live, excised, lymph nodes, we used mice bearing a combination of XCR1-Venus(*18*), CD11c-mCherry(*19*) and MacBlue(*20*) transgenes. These alleles were chosen as they largely discriminate key populations under study here, namely: Venus^+^mCherry^+^ = cDC1, mCherry^+^ = cDC2, CFP^+^ = monocytes and a fraction of mCherry^+^CFP^+^ = resident cDC2 (**Fig. 3A, Fig. S7**). These cells are in constant, albeit slow, motion within the lymph node; we measured the duration of engagements, defined as the edge of the fluorescence of one cell type within 1μm of another (**Fig. 3A**)(*21*). We then plotted the lifetimes of the interactions (**Fig. 3B**). The time of interaction between populations was best modeled by a two-phase exponential decay. The first phase for all contact pairs had a T^1/2^ of <60s, likely representing cells passing one another (**Fig. 3B**). The second phase only occurred in about 20% of the contacts between two DC and demonstrated a much longer T^1/2^ of approximately 240s with durations that could extend beyond 10 minutes (**Fig. 3B-C**). This suggested that, at times, DC might engage in more substantial communication with one another. Of note, only ∼5% of monocyte:DC contacts endured into the second phase of decay (**Fig. 3B-C**).

**Fig. 3.**
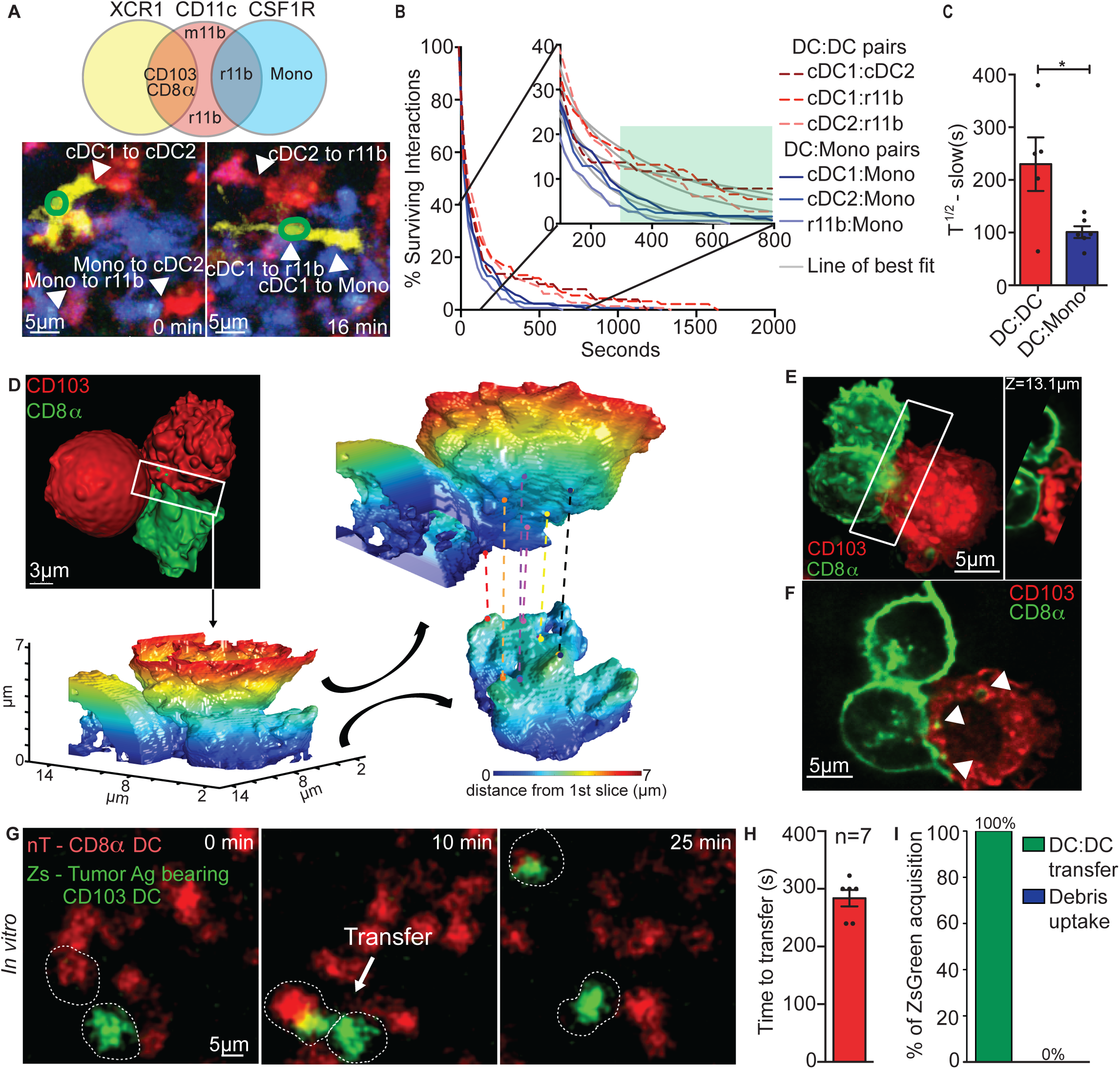
Dendritic cells form stable, heterotypic interactions and pass tumor-derived antigen. (**A**) Venn diagram showing triple reporter (XCR1-Venus, CD11c-mCherry, CSF1R-CFP) gene expression for various myeloid cells in the tumor draining lymph node (tdLN, top). Representative multiphoton microscopy images of tdLN explants of B16ZsGreen tumor-bearing, triple reporter mice. Images are a timecourse showing a ZsGreen-containing XCR1-Venus^+^ cell interacting with other myeloid cells in the tdLN and other unloaded myeloid cells interacting with each other. Interactions indicated by labeled arrows. (**B**) Interaction times of myeloid cells within the tdLN. Inset is zoomed in view to show detail of differential interaction times. Grey line is line of best fit assuming 2-phase exponential decay. Green box denotes second phase of decay. (**C**) Quantification of interaction time half-life (T^1/2^) for dendritic cell (DC) to DC interactions and of DC to monocyte (Mono). Data are representative of 3 experiments. Plot shows mean T^1/2^of second phase decay +/−SEM. * < 0.05. (**D**) Lattice light sheet (LLS) image of interaction interface of sorted CD103 DC from membrane-tdTomato (mT) mouse and CD8α DC from membrane-GFP (mG) mouse. White box shows interface location used for contact analysis. Heatmap represents distance from reference z-slice (dark blue = 0). Interface is opened (black arrows) and rotated to show surface points of juxtaposed sites indicated with dots connected with color-coded dotted line. (**E**) LLS image showing membrane proximity of mG^+^CD8α^+^resident DC (green) and mT^+^CD103^+^ DC (red). White box highlights interaction surface shown as z-slice (right). Scale bar = 5µm. (**F**) Single z-slice of cells in E highlighting points of membrane exchange (white arrrows). Scale bar = 5µm. (**G**) Live, confocal microscopy showing ZsGreen^+^ CD103^+^DC (green) sorted from B16ZsGreen tdLN and a nuclear-tdTomato (nT) CD8α^+^ resident DC (red) sorted from steady state nTnG mouse. 2 individual cells are marked with dotted lines throughout and transfer event is shown in the middle panel. Scale bar = 5µm. (**H**) Quantification of the average contact time between cells preceding antigen transfer *in vitro*. Data obtained from imaging described in G. (**I**) Frequency of ZsGreen acquisition based on mode of uptake (debris uptake versus DC:DC transfer) quantified from observation of imaging timecourses in G. n = 6.

To examine the details of such DC-DC interactions, we used high-resolution, real-time, LLS *in vitro*(*21*) using tdLN DC from mice in which the membranes were genetically marked with fluorophores. This revealed a consistent landscape of co-conformations at the DC-DC interface with some forming a stable contact (**Fig. 3D-F, Fig. S8A-B, Movie S2**). In 6 out of 16 synapses analyzed, we observed only minimal membrane engagements with little surface complementarity (**Fig. S8A**). However, 10 out of 16 synapses demonstrated engagements exemplified in **Fig. 3D, Movie S2**, and **Fig. S8B**. When we separated and rotated these synapses, we found that both cells faced one another ‘cup-on-cup’ (i.e. each membrane concave in the center of the contact). In one rare example, we also found a finger-like projection protruding into the cup (**Fig. S8B**). The sum of the LLS studies show that when DC contact one another, they initially probe with veils of membrane but can further engage in more persistent close membrane-membrane juxtapositions (**Fig. 3D-F, Fig. S8, Movie S2-4**). Notably, the active synapses also showed evidence of exchange of both vesicular material and membranes (arrows in **Fig. 3F, Movie S5, Fig. S9**).

We thus hypothesized that these synapses might represent the structure for cell-to-cell tumor antigen-containing vesicle transfer. To investigate the outcome of these interactions we first imaged the *in vitro* mancala assay from **Fig. 2E** (**Fig. 3G**), now using a conventional confocal microscope with a wider field of view. We took time-lapse movies of a single type of transfer—from the typically dominant migratory CD103^+^ cDC1, into the resident CD8α^+^ population. Antigen loaded CD103^+^ DC were observed forming contacts and then transferring a portion of their ZsGreen cargo into the recipient, in the absence of a secreted intermediate (**Fig. 3G, Movie S6**). Antigen transfer occurred on timescales consistent with the 2^nd^ phase observed *in vivo*, typically occurring within approximately 300s after contact formation (**Fig. 3H**). Transfer overall was rare, occurring in <1% of contacts in the typical 8-10 hours of imaging, but was observed in all pairs of migratory donor and a resident recipient (data not shown). We also observed ZsGreen-loaded CD103^+^ DC passing vesicles sequentially to multiple CD8α^+^ DC (**Movie S7**) which shows the process can be repeated and that a cell can pass antigen while remaining viable both before and after the passing occurs. In support of synaptic transfer as the dominant mechanism of handoff, antigen acquisition by recipient DC was always observed in the context of contact between cells and never observed as uptake of cell-free/apoptotic debris, in over 48 hours of total imaging (**Fig 3I**).

Motivated to examine antigen transfer *in vivo*, we performed multiphoton microscopy in the interfollicular T cell zone of tdLN of XCR1-Venus; CD11c-Cherry; MacBlue mice bearing B16ZsGreen tumors. An example, shown in **Fig. 4A** (**Movie S8**), shows a cDC2 (CD11c-Cherry^+^) transfering ZsGreen to a cDC1 (XCR1-Venus^+^). As *in vitro*, overall frequencies of transfer in a typical 30 minute movie was rare, but approximately 75% of the imaging experiments resulted in at least one detectable transfer event, which always occurred at a contact and never via a DC-free exosome or apoptotic body (**Fig. 4B**).

**Fig. 4.**
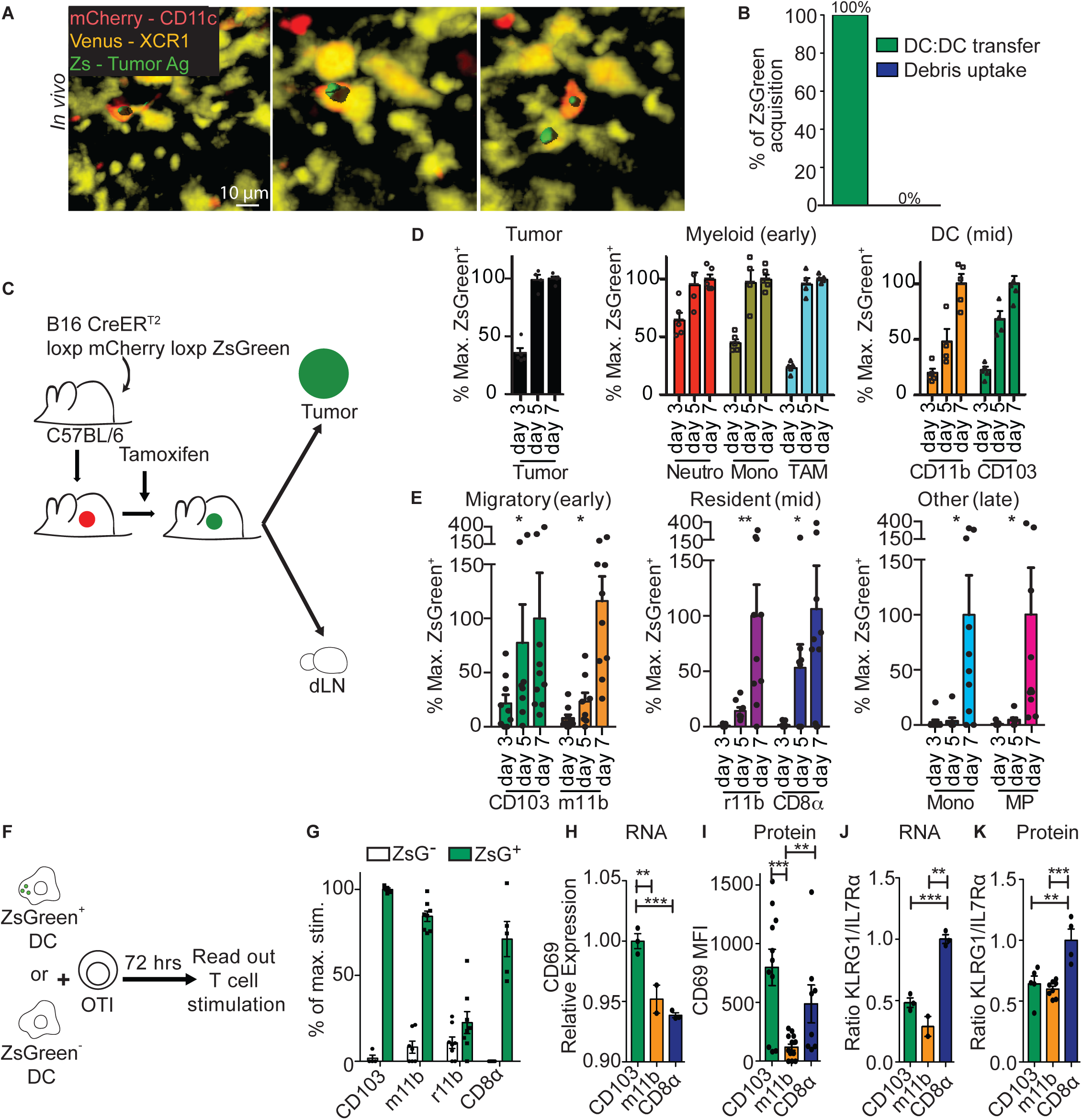
Temporal tracking of tumor antigen flow *in vivo* and consequential T cell priming characteristics. (**A**) Live, multiphoton microscopy of tumor-draining lymph nodes (tdLN) from B16ZsGreen, XCR1-Venus; CD11c-mCherry; MacBlue mice. For clarity, only Venus and mCherry channels shown. ZsGreen^+^ vesicle is surfaced (green); timecourse shown. Scale bar = 10µm. (**B**) Frequency of ZsGreen acquisition based on mode of uptake (debris uptake versus DC:DC transfer) quantified from imaging timecourses in a. n = 3. (**C**) Method for inducible-ZsGreen B16 tumor experiments in D-E. (**D-E**) ZsGreen accumulation within cells of the tumor (D) and tdLN (E) following induced expression of ZsGreen. Days indicate initiation of tamoxifen treatment prior to sacrifice. % Max. ZsGreen^+^ calculated as mean frequency of ZsGreen^+^ cells normalized to the average maximum % of ZsGreen accumulation (set to 100%) for each cell type analyzed. Representative of 3 independent experiments. (**F**) Experimental method for *in vitro* T cell stimulation assay in G. (**G**) % of maximum CD8^+^, OTI, T cell stimulation following 72hrs of coculture with different DC types that are either ZsGreen^+^ (green bars) or ZsGreen^−^ (white bars). Setup shown in F. Plot shows mean % of maximum OTI stimulation as assessed by dilution of eFluor670 dye. Error bars represent +/−SEM. Data is 3 independent experiments combined. (**H**) Relative expression of CD69 in stimulated T cells as determined by RNA sequencing. Error bars represent +/−SEM. Data show 2 experimental replicates for m11b DC and 3 replicates for CD103 and CD8α DC. (**I**) Flow cytometric mean fluorescence intensity (MFI) of CD69 staining on OTI T cells stimulated with the different DC types. Error bars are +/−SEM. Representative of 3 independent experiments. (**J**) Ratio of KLRG1/IL7Rα RNA expression levels for OTI T cells stimulated with different DC types. Error bars represent +/−SEM. Data show 2 experimental replicates for m11b DC and 3 replicates for CD103 and CD8α DC. (**K**) Normalized ratio of KLRG1 MFI to IL7Rα MFI in OTI T cells stimulated with the different DC types. Error bars are +/−SEM. Representative of 3 independent experiments. In all graphs **p < 0.01, ***p < 0.001.

Direct cell contacts would predict a sequential filling of antigens into various myeloid populations as one directly hands off to others, as opposed to all compartments filling at the same rate. To test this *in vivo*, we developed a method to pulse tumor material into antigen presentation by generating a B16F10 line initially expressing mCherry which could be induced to express ZsGreen by virtue of the expression of tamoxifen regulated Cre (**Fig. 4C**). This tumor line was injected into mice, which were then gavaged with tamoxifen at staggered time points prior to harvest at day 14 post-injection. Tumor and tdLN were then examined for ZsGreen fluorescence (**Fig. 4C**). In the tumor, cells acquired ZsGreen with distinct kinetics: neutrophils, monocytes and macrophages became maximally ZsGreen^+^ in near-concert with the tumor itself, maximally by day 5 (**Fig. 4D**). Tumor-resident DC loading kinetics were more protracted and maximal loading occurred by day 7 (**Fig. 4D**). Although there was greater variability between mice in the tdLN, there was a clear hierarchy wherein only migratory DC populations were significantly loaded at day 3 and continued to increase thereafter (**Fig. 4E**). In contrast, no loading was observed for resident DC until day 5. Finally, monocytes and microparticles (MP) containing ZsGreen were only detected at day 7 (**Fig. 4E**). This result is consistent with older work in vaccination models(*22*) that demonstrated a need for cell-dependent delivery of material to lymph nodes but here is extended to tumor antigens with vesicular hand-off specific myeloid subpopulation kinetics.

To examine whether ZsGreen vesicle passage is required for DC to obtain significant ligands for T cells, we generated B16F10 cells expressing ZsGreen fused to the OTI and OTII OVA peptides (B16ZsGreen-minOVA). This allowed for initial flow-sorting of otherwise similar DC into ZsGreen^+^ versus ZsGreen^−^ subsets and then permitted us to test which population could stimulate T cells (**Fig. 4F-G and Fig. S10A-C**). This revealed that only DC marked by detectable ZsGreen levels could stimulate OTI T cells, regardless of subset (**Fig. 4G, Fig. S10B**). If other methods of antigen-transfer were dominant, we could expect that both populations would stimulate similarly. We may conclude that other mechanisms that do not accompany such transfer are relatively ineffective sources of ligands. While it is possible that pre-loaded pMHC is acquired concurrently with bolus antigen, work by others has shown cross-dressed DCs are only effective in stimulating resting memory T cells, and are unable to activate a naïve population(*17*). It may thus be that low levels of membrane-only transfer is sufficient to trigger highly activated T cells but would likely have less importance for naïve T cell priming.

The ability to track antigens into specific recipient DC has further consequences for understanding diversification of T cell response. Previous reports from our lab and others, that did not isolate the exact cells bearing bolus antigen, had concluded that migratory CD103^+^ cDC1 were unique in their ability to profoundly stimulate anti-tumor CD8^+^ T cells(*1, 14*). However, when ZsGreen antigen-bearing cells were selectively purified, we now also observed stimulatory capacity for resident CD8α^+^ DC and migratory CD11b^+^ DC (**Fig. 4G**). (CD4^+^ OTII cells were stimulated most efficiently by ZsGreen^+^ migratory CD11b^+^ DC as previously reported(*23*)(**Fig. S10C)**). Beyond revealing the undocumented CD8^+^ T cell stimulatory capacity of migratory CD11b^+^ and resident CD8α^+^ DC, the selection of antigen-bearing DC also now allowed direct *ex vivo* analysis of the induction of distinct effector programs by DC subsets. It has previously been reported that different DC can drive T cells to differentiate in distinct ways during infection(*24*) and following acquisition of dying self-cells(*25*), but heretofore, has not been shown in the tdLN. Comparing RNA expression from T cells following stimulation with the three proliferation-driving DC subsets demonstrated that the ZsGreen vesicle^+^, CD8α^+^ DC, drove the most transcriptionally distinct outcome, with 291 genes consistently differentially regulated by CD8α^+^ DC compared to both CD11b^+^ DC and CD103^+^ DC (**Extended Data Fig. 8d-e**). These genes were enriched in signature GO terms relevant to immune function comprising genes involved in interferon responses, cell division and T cell mediated cytotoxicity (**Fig. S10D, Fig. S11**). By comparing the gene expression profiles to previously defined molecular signatures (*26, 27*), we find that cells stimulated via resident CD8α^+^ DC show reduced responses to IFNγ pathway induction and memory induction and increased modules associated with short term effectors (**Fig. S12**). Specific distinctions in RNA were confirmed at the protein level, demonstrating specific decreases in PD1 expression following priming via migratory CD11b^+^ DC (**Fig. S10F-G**), increased CD69 expression when stimulated by CD103^+^ DC (**Fig. 4H-I**) and the combination of decreased CD127 and increased KLRG1 (**Fig S10G**, as a ratio in **Fig 4J-K**) when triggered by antigen-positive CD8α^+^ DC. The latter is once again significant for antigen handoff since a skewed ratio of these genes is associated with the formation of short-term effector subsets in other systems(*28*) and thus tumor loading of these cells would skew away from memory phenotypes, which are thought to be more capable of tumor rejection(*29, 30*).

Studies of tdLNs in human melanoma patients have observed vesicles, similar to those we described here, inside myeloid populations termed ‘melanophages’(*31, 32*). While other mechanisms, not observed in this study, may provide additional antigens to T cells, our data clearly supports that specific and contained vesicles of antigen are a robust mechanism for antigen handoff, leading to tdLN priming and potentially contributing to tumor tolerance. The ‘mancala’ handoff mechanism moves this field well beyond ‘trogocytosis’ (*33*), defined as the nibbling of one cell by another, insofar as this vesicle delivery is the very deliberate transfer of a discrete ‘packet’ of material which is highly enriched in material from an original donor. And while “cross-dressing” defined as movement of intact peptide-MHC complexes and important for memory T cell activation (*17*) may happen independently, contemporaneously or subsequent to the vesicular exchanges we’ve identified here, our results positively correlate cells containing these vesicles with the ability to stimulate naïve T cells. It may be that the vesicles transferred carry information, beyond the antigens themselves, that are instructive to myeloid cells and the synapses described in this study presumably provide additional opportunities for the immune system to self-align. Further studies will need to define how these vesicles are initially programmed at the tumor or elsewhere; options including via nibbling of live cells, processing of dead debris or ingestion of pre-formed exosomes. The possible role of these vesicles for transfer of non-self commensals and pathogen-derived will also be important. Referencing the two-signal model of T cell activation—wherein signal 1 (TCR-pMHC signaling) is complemented by signal 2 (CD28, and opposed by CTLA-4 and PD1 checkpoints)— manipulation of this transfer biology, affecting which cell type presents signal 1 and how much signal 1 it presents should represent an excellent orthogonal strategy to checkpoint blockade therapies, to license or augment therapeutic benefits.

## Materials and Methods

### Mice

Mice were housed and bred under specific pathogen-free conditions at the University of California, San Francisco Laboratory Animal Research Center and all experiments conformed to ethical principles and guidelines approved by the UCSF Institutional Animal Care and Use Committee. C57BL/6 mice were purchased from Jackson Laboratory or bred in house, and unless otherwise noted animals used were male between 6–8 weeks of age. C57BL/6 were used for all ectopic tumor studies. *Ccr7*-deficient C57BL/6 mice, purchased from Jackson Laboratory, were used for modulation of tumor-associated DC migration studies(*34*). The mTmG reporter strain (*35*) was maintained both as a single transgenic and also crossed with a βactin-Cre line (Jackson Laboratory) to achieve mG mice. nTnG mice were purchased from Jackson Laboratory. *Ccr2*-deficient C57/BL6 mice were purchased from Jackson Laboratory. XCR1-Venus (*18*) mice were crossed with CD11c-mCherry (Jackson Laboratory) and MacBlue (Jackson Laboratory) reporter mice to achieve the triple reporter mouse line. OTI and OTII mice were purchased from Jackson Laboratory and then crossed with CD45.1+ congenic mice for use as T cell donors in the T cell stimulation assays. CD45.1 mice were obtained from Jackson Laboratory.

### Cell lines and ectopic tumor injections

Briefly, adherent cells were cultured at 37**°**C with 5% CO_2_ in DMEM (GIBCO) plus 10% heat-inactivated FCS with penicillin, streptomycin and L-glutamate on tissue culture-treated plastic plates and split every other day. B16ZsGreen, B16 melanoma parental cells were genetically engineered to stably express ZsGreen using viral transduction with a ZsGreen construct. Inducible B16ZsGreen, B16 melanoma parental cells were genetically engineered to stably express both a Thy1.1-CreER^T2^ construct and a loxP-mCherry-loxP-ZsGreen construct using viral transduction of both expression vectors and selection for stable integrants using an Aria II cell sorter and detection of mCherry and Thy1.1 co-expressing cells. B16zsGreen-minOVA, B16 melanoma parental cells were genetically engineered to stably express ZsGreen-minOVA using viral transduction with a ZsGreen-minOVA construct. For ectopic tumor injections, cells were grown to confluency, harvested and washed 2 times with PBS, mixed at a 1:1 ratio with growth factor-reduced Matrigel Matrix (BD Biosciences) in a final injection volume of 50ul. Two hundred thousand tumor cells were injected subcutaneously in the right and left flanks of mice and allowed to grow for 14 days before harvest of tumor and lymph node.

### Tissue digest and DC isolation

LN were dissected from tumor bearing or WT mice, and cleaned of fat. For tumors, inguinal and axillary LN were taken as tumor draining. LN were digested as previously described (1). In brief, LN were pierced and torn with sharp forceps in 24-well plates and incubated for 15 min at 37° C in 1 ml digestion buffer (100 U/ml collagenase A (Roche), 500 U/ml collagenase D (Roche), and 20 ug/ml DNAse I (Roche) in RPMI-1640 (GIBCO)). After the first 15-minute incubation, cells was pipetted up and down repeatedly, and, then returned for a second 15-minute incubation at 37 ° C. After digestion, LN were washed with RPMI-1640 (GIBCO) plus 10 % FCS and filtered through 70 um Nytex filters before staining for flow cytometry. When sorting from LN, cells were stained with biotin-conjugated anti-CD2 (clone RM2-5, BioLegend) antibody and negative selection was performed by using an EasySep Biotin Selection Kit (Stemcell Technologies) following manufacturers instructions, before staining. For tumor digests, tumor were isolated and minced prior to 30 minute incubation in 6 mL digestion mix (same as LN above) on a shaker at 37 ° C.

### Flow cytometry and antibody clones

For surface staining, cells were incubated with anti-Fc receptor antibody (clone 2.4G2, UCSF Hybridoma Core) and then stained with antibodies in PBS + 2 % FCS for 30 min on ice. Viability was assessed by staining with fixable Live/Dead Zombie (BioLegend) or DAPI. Flow cytometry was performed on a BD Fortessa instrument. Analysis of flow cytometry data was done using FlowJo (Treestar) software. Cell sorting was performed using a BD FACS Aria II or BD FACS Aria II Fusion.

Antibodies were used against the following targets: CD11b Brilliant Violet 605 (clone M1/70, BioLegend), CD205 PerCP-Cy5.5 (clone NLDC-145, BioLegend), CD103 APC (clone 2E7, BioLegend), CD8α PE-Cy7 or PerCP-Cy5.5 (clone 53-6.7, BioLegend), CD150 APC-Fire 750 (clone TC15-12F12.2, Biolegend), CD11c Brilliant Violet 510 or Brilliant Violet 650 (clone N418, BioLegend), Ly6C Brilliant Violet 711 (clone HK1.4, BioLegend), Ly6G Brilliant Violet 785 (clone 1A8, BioLegend), B220/CD45R Brilliant Violet 785 (clone RA3-6B2, BioLegend), I-A/I-E Brilliant Violet 421 (clone M5/114.15.2, BioLegend), biotin conjugated anti-F4/80 (clone BM8, eBioscience) in conjunction with streptavidin Brilliant Violet 510 (BioLegend), CD24 PE-Cy7 (clone M1/69, BioLegend), CD45 Alexa 700 (clone 30-F11, BioLegend), CD90.2 Brilliant Violet 785 (clone 30-H12, BioLegend). Annexin V staining was performed using an APC Annexin V staining kit (BioLegend) according to manufacturer’s instructions.

### Imaging

Confocal Imaging: DC populations were sorted (based on the gating strategy in Figure S1A) using a FACS Aria II flow cytometer from tumor-draining LN and plated onto fibronectin-coated glass slides (ZsGreen puncta imaging) or in wells of a fibronectin-coated 384-well plate (live, *in vitro* antigen transfer assay) and then imaged on a Leica SP5 laser scanning confocal microscope. For puncta staining, cells were fixed (0.05 M phosphate buffer containing 0.1 M L-lysine (pH 7.4), 2 mg/ml NaIO4 and 1% PFA), blocked and permeabilized (1% normal mouse serum, 1% bovine serum albumin and 0.3% Triton X-100) then stained with anti-gp100 antibody (clone HMB45, BioLegend) used at 1:30 followed by an anti-IgG2a secondary conjugated to Alexa555 (Invitrogen) at a concentration of 1:250. Anti-tyrosinase antibody (clone TA99, BioLegend) was used at 1:50 followed by an anti-IgG1 secondary conjugated to Alexa647 (Invitrogen) at a concentration of 1:250. The inverted microscope system is encased within an incubator for live-cell imaging overnight. For whole tissue imaging of LN and tumor, methods for fixing and clearing tissues were performed as previously described(*36*). Tissues were then imaged using a Nikon A1R laser scanning confocal microscope using NIS-Elements software. Data analysis was performed using the Imaris software suite (Bitplane).

#### LLS

LLS imaging was performed in a manner previously described(*21*). Briefly, 5 mm diameter round coverslips were cleaned by a plasma cleaner, and coated with 2 μg/ml fibronectin in PBS at 37°C for 1 hour before use. Sorted DC were dropped onto the coverslip and incubated at 37°C, 5% CO_2_ for 20–30 min. The sample was then loaded into the previously conditioned sample bath and secured. Imaging was performed with a 488-nm, 560-nm, or 642-nm laser (MPBC, Canada) dependent upon sample labeling. Exposure time was 10 ms per frame leading to a temporal resolution of 4.5 s. For cell surface labeling of CD45.1+ recipient DC, anti-CD45 (30-F11, BioLegend) directly conjugated to Alexa647 was used at a concentration of 1:200 for 20min prior to seeding coverslip. Image renderings were created using ChimeraX software(*37*).

#### Two Photon Microscopy

Intravital imaging was performed using a custom-built two-photon setup equipped with two infrared lasers (MaiTai: Spectra Physics, Chameleon: Coherent). The MaiTai laser was tuned to 800 nm and the Chameleon laser excitation was tuned to 950 nm. Emitted light was detected using a 25× 1.2 NA water lens (Zeiss) coupled to a 6-color detector array (custom; utilizing Hamamatsu H9433MOD detectors). Emission filters used were: violet detector 417/60, blue 475/23, green 510/42, yellow 542/27, red 607/70, far red 675/67. The microscope was controlled by the MicroManager software suite, z-stack images were acquired with 4-fold averaging and z-depths of 3 µm. Data analysis was performed using the Imaris software suite (Bitplane). To characterize contact parameters DCs, tracks and surface of different DCs were generated, and the dwell time of interaction between surfaces was analyzed as previously described (*38*): DC–DC interaction was defined as the association at 1 μm or less of a given DC cell surface with another DC surface.

### LN explant imaging of *in vivo* antigen transfer

Imaging was performed as previously described (*38*). Inguinal and axillary LN were removed, cleaned of fat, and immobilized on a plastic coverslip with the hilum facing away from the objective. LN were imaged in 30-minute intervals on a two-photon microscope as per above.

### *In vitro* antigen transfer assays

Recipient nT^+^ DC were isolated and sorted as described above from LN of nTnG mice. ZsGreen^+^ donor DC are sorted from the tumor-draining LN of B16ZsGreen tumor-bearing mice. Then cells are plated in complete media (RPMI-1640 plus 10% heat-inactivated FCS with penicillin, streptomycin, L-glutamate, non-essential amino acids) supplemented with 7.5 ng/mL GMCSF for 16hrs as either individual, non-competitive transfer assays or competitive, equimolar transfer assays in 96 well V-bottom tissue culture-treated plates prior to staining and analysis by flow cytometry.

Non-competitive antigen transfer assay: sorted cells are plated for individual recipient transfer where 6000 total cells per well are plated at a 1 nT^+^ recipient:1 ZsGreen^+^ donor DC ratio.

Competitive antigen transfer assay: sorted cells are plated with all 4 recipient DC types combined together maintaining a 1:1 ratio of donor:recipient and a total cell number of 6000 cells/well.

Transwell antigen transfer assay: sorted cells are plated in the same way as the equimolar antigen transfer assay with the addition of a 3 μm pore size transwell insert which separates donors from recipients. Donors are plated in the top of the insert and recipients were plated in the bottom.

Inhibitors of various cellular processes were used where indicated and added at the time of cell plating: GDC0941 at a concentration of 10μM, VPS34-IN1 at a concentration of 1μM, and zVAD at a concentration of 20μM.

### T cell stimulation assays

OT1 and OTII T cells were isolated from LN of TCR transgenic mice using either a CD8 or CD4 StemSep enrichment kit (STEMCELL Technologies), respectively. DC were obtained through sorting from the dLN of B16zsGreenminOVA bearing mice as described above for B16zsGreen tumor dLN. DC were sorted as either ZsGreen^+^ or ZsGreen^−^ for each of the 4 DC subsets and subsequently used for T cell stimulation. Stimulation assay was performed as previous described (1). Dilution of cell permeable dye eFluor670 (eBioscience) and expression of CD44 (IM7, BioLegend) were used as indicators of T cell stimulation.

### RNA-sequencing

STAR 2.4.2a was used to align reads to the Mus Musculus genome, version GRCm38.78. Only mapped reads uniquely assigned to the mouse genome were used for differential expression testing. These were imported into R and then converted to normalized read counts with DESeq2. QC plots were created from the counts generated by DESeq2’s variance Stabilizing Transformation (with blind=True). Differential Expression was performed using DESeq2, and significant genes were filtered by a q-value (False Discovery Rate) threshold of 0.05, and a p-value threshold of 0.05. pheatmap was used for correlation heatmaps, and heatmap.2 for the QC heatmap. Additionally, gene ontology (GO) analysis was performed using the publically available resource from the GO Consortium and the generated RNA-sequencing data.

### Statistical analysis

Statistical analyses were performed using GraphPad Prism software. Unless specifically noted, all data are representative of >3 separate experiments. Error bars represent standard error of the mean (S.E.M.) calculated using Prism, and are derived from triplicate experimental conditions. Specific statistical tests used were paired and unpaired t-tests and p values <0.05 were considered statistically significant. GO analysis was performed using the RNA-sequencing data

## Acknowledgments

We thank D. Hume for providing MacBlue mice; T. Kaisho for providing XCR1-Venus mice; J. Cyster for providing us with CCR2-knockout mice. We would also like to thank A. Gérard for her initial work in developing explanted LN imaging. K. Corbin for early technical assistance in this project. All members of the Krummel laboratory, BIDC, and A. Denton for discussion, support, and guidance while developing this work.

## Author contributions

E.W.R and M.K.R. designed and conducted most of the experiments, data analysis, and drafted the manuscript; E.C. performed LLS data analysis; A.M.M. performed the CCR2 knockout mice experiments and triple reporter LN flow cytometry; K.M., and C.B. assisted with the LLS; D.N. generated the B16 CreER^T2^ loxp-mCherry-loxp-ZsGreen B16ZsGreen cell line; N.S. and M.B. discussed data and project direction; M.F.K. designed experiments, interpreted data, and with other authors, developed the completed manuscript.

## Data and materials availability

All data included in this study is available from authors upon reasonable request. RNA sequencing data has been submitted to GEO and can be found using the accession number GSE128980.

