## Supplemental Data for "Tumors Exploit Dedicated Intracellular Vesicles to Program T cell Responses"

**SUPPLEMENTARY MATERIALS**

**Figures S1-S12**

**Captions for Movies S1 to S8:**

**Movie S1: 3D rendering of lymph node resident CD11b$^+$ dendritic cell harboring tumor-derived ZsGreen as puncta**

Lattice light sheet imaging of sorted dendritic cell (DC) from B16ZsGreen tdLN. Three dimensional rendering of cell shown as z-slice in **Fig. 1F**. Representative image. Scale bar = 2µm.

**Movie S2: Analysis of tight synaptic engagement between CD103$^+$ DC and CD8α$^+$ DC**

Representative lattice light sheet, heatmap-surfaced, image examining the contact interface between CD103$^+$ DC (top) and CD8α$^+$ DC (bottom) from the static image shown in **Fig. 3D.** Synapse has been opened and is rotating to show topology of the membrane at the contact. Juxtaposed sites indicated with dots connected with color-coded dotted line.

**Movie S3: Contact formation between mT$^+$ CD103$^+$ DC and mG$^+$ CD8α$^+$ DC**

Representative lattice light sheet live-imaging of mT$^+$ migratory CD103$^+$ DC (red) and mG$^+$ LN resident CD8α$^+$ DC (green) sorted from steady state LN. Data are representative of 2 independent experiments. Scale bar and frame rate as indicated.

**Movie S4: Z series through contact between mT$^+$ CD103$^+$ DC and mG$^+$ CD8α$^+$ DC**

Representative lattice light sheet live-imaging frame with z projection showing close membrane proximity of mG$^+$ LN resident CD8α$^+$ DC (green) and mT$^+$ migratory CD103$^+$ DC (red).  Cells were sorted from steady state LN of mT or mG mice. White box drawn to highlight interaction surface shown as a z-series (right). Data are representative of 2 independent experiments. Scale bar as indicated.

**Movie S5: Membrane exchange between mT$^+$ CD103$^+$ DC and mG$^+$ CD8α$^+$ DC**

Representative lattice light sheet live-imaging of single z plane showing membrane exchange between mG$^+$ LN resident CD8α$^+$ DC (green) and mT$^+$ migratory CD103$^+$ DC (red). Cells were sorted from steady state LN of mT or mG mice. Red arrows indicated CD103$^+$ red membrane transferred to CD8α$^+$ mG$^+$ DC. Green arrows indicate green membrane transferred to mT$^+$ CD103$^+$ DC. Data are representative of 2 independent experiments. Scale bar and frame rate as indicated.

**Movie S6: Transfer from ZsGreen$^+$ CD103$^+$ DC to nT$^+$ CD8α$^+$ DC**

Live confocal microscopy imaging showing ZsGreen$^+$ CD103$^+$ DC (green) sorted from tumor draining LN and nT$^+$ CD8α$^+$ LN resident DC (red) sorted from steady state nTnG mouse. Transfer event is indicated with an arrow. Movie is shown with and without differential interference contrast (DIC). Dotted lines indicate cell membrane. Data are representative of 5 independent experiments. Scale bar and frame rate as indicated.

**Movie S7: Sequential transfer from ZsGreen$^+$ CD103$^+$ DC to nT$^+$ CD8α$^+$ DC**

Live confocal microscopy imaging showing ZsGreen$^+$ CD103$^+$ DC (green) sorted from tumor draining LN and nT$^+$ CD8α$^+$ LN resident DC (red) sorted from steady state nTnG mouse. Two transfer events followed by full abscission are indicated with arrows. Data are representative of 5 independent experiments. Scale bar and frame rate as indicated.

**Movie S8: Transfer of ZsGreen in ex vivo LN from XCR1-Venus, CD11c-mCherry, MacBlue mouse**

Representative mulitphoton confocal live imaging of tumor draining LN explants from B16ZsGreen tumor-bearing XCR1-Venus, CD11c-mCherry, MacBlue mice. Time course is first shown with ZsGreen vesicle surfaced in green followed by same time course without surfacing. MacBlue channel is not shown for clarity. Data are representative of 3 independent experiments. Scale bar and frame rate as indicated.

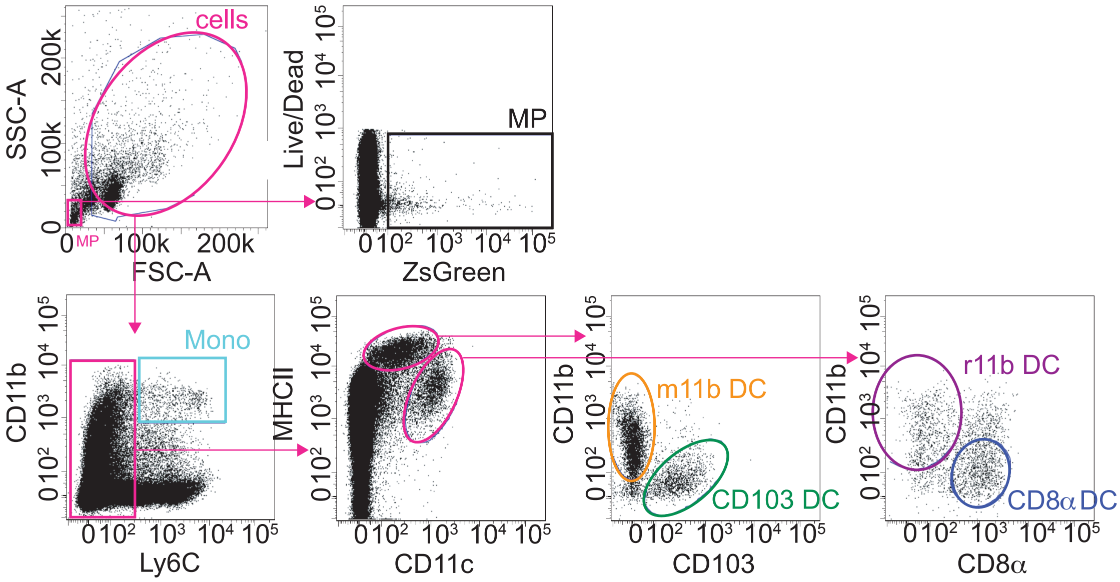


**Fig. S1 Gating strategy for identification of myeloid cell types in lymph nodes**
Flow cytometric gating strategy for the delineation of the B16ZsGreen melanoma tumor-draining lymph node (tdLN) myeloid compartment. Combined inguinal and axillary tdLN. Progressive gating identifies LN microparticles (MP, black), resident $CD8\alpha^+$ dendritic cells (DC, blue), migratory $CD103^+$ DC (green), and resident (purple) and migratory (orange) $CD11b^+$ DC subsets.

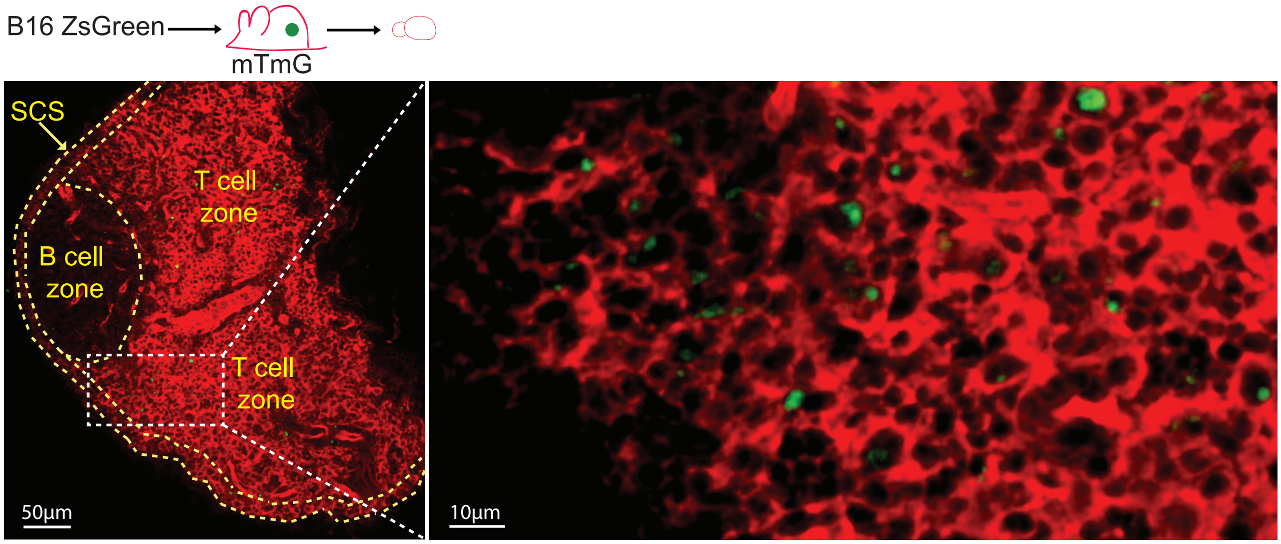


**Fig. S2 ZsGreen distribution in the tumor-draining lymph node**
Schematic showing experimental setup for tumor-draining lymph node (tdLN) imaging (top). Image from cleared B16ZsGreen, tdLN demonstrating overall ZsGreen distribution at lower magnification (left, scale bar = 50μm) and higher magnification (right, scale bar = 10μm). LN regions are outlined and labeled in yellow. Higher magnification image is zoomed in from white box. Representative images.

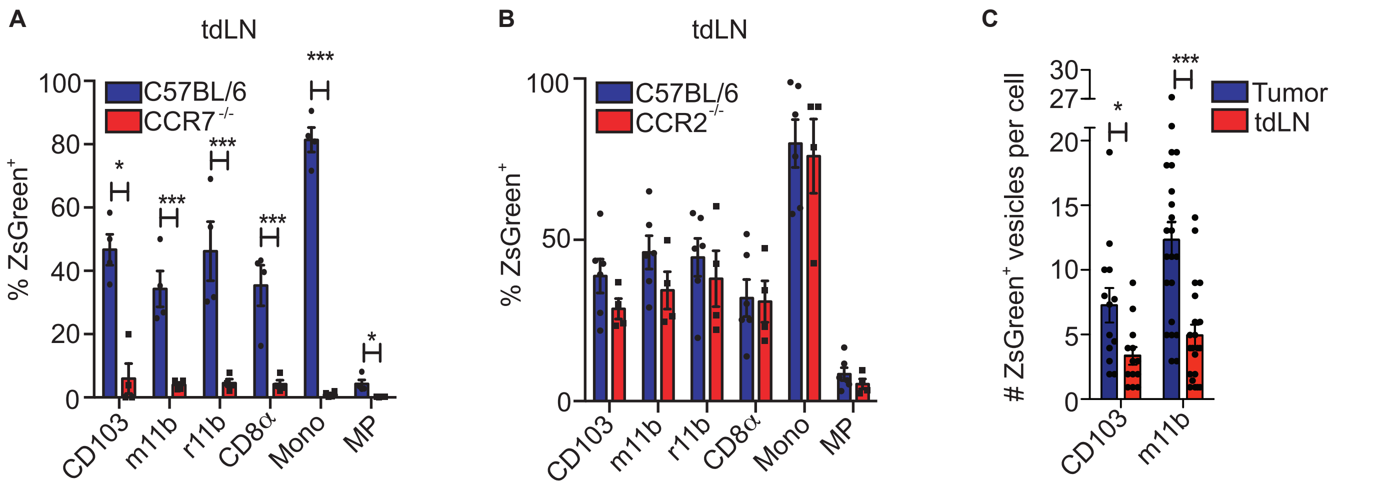


**Fig. S3: Trafficking of tumor-derived antigen is dependent on myeloid cell migration and not mediated through monocyte migration to the lymph node**

(**A**) Quantification of tumor-derived ZsGreen within myeloid cells or microparticles (MP) of tumor draining LN (tdLN) from C57BL6 wildtype or CCR7 $^{-/-}$ mice. Plot shows mean frequency +/- SEM. Data are representative of 3 independent experiments. migratory CD11b DC (m11b), resident CD11b DC (r11b). (**B**) Quantification of tumor-derived ZsGreen within myeloid cells or microparticles (MP) of the tdLN in C57BL/6, wildtype mice or CCR2 $^{-/-}$ mice. Plot shows mean frequency +/- SEM. Data are representative of 2 independent experiments. (**C**) Quantification of mean ZsGreen $^{+}$ vesicle count per cell in migratory DC subsets (CD103, m11b) in the tumor and tdLN. Plot shows mean frequency +/- SEM. *p < 0.05, **p < 0.01, ***p < 0.001.

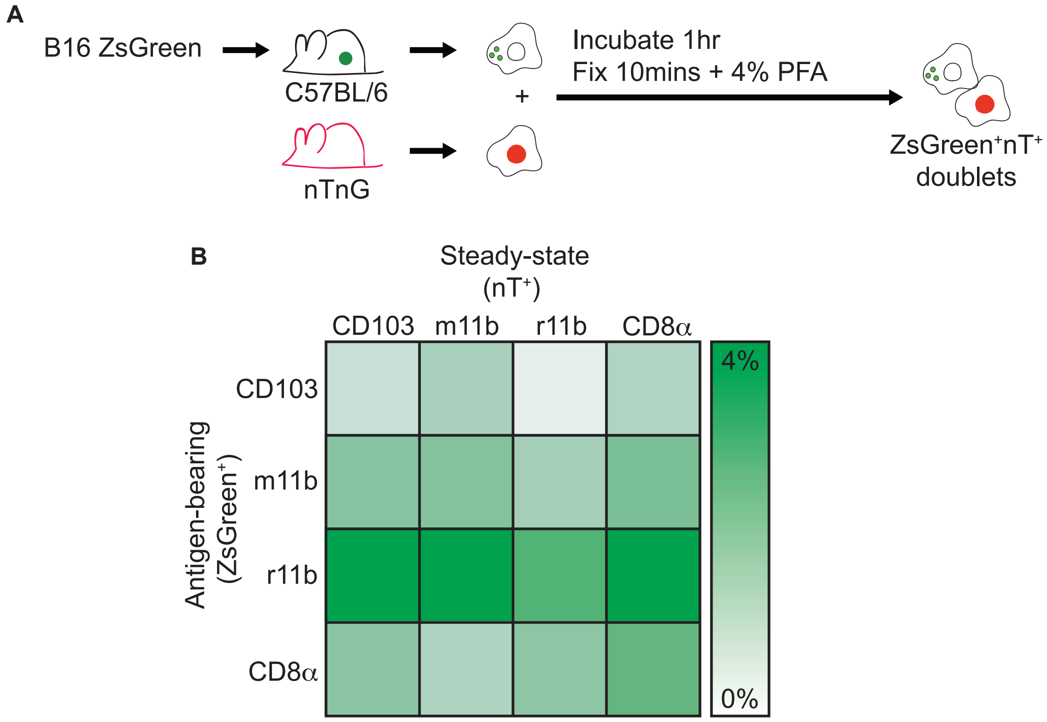


**Fig. S4: Dendritic cells from tumor-draining lymph nodes form lasting interactions *in vitro***
(**A**) Schematic showing experimental method of *in vitro* dendritic cell (DC) coupling assay. Tumor antigen-loaded DC (ZsGreen$^+$) are sorted from B16ZsGreen tumor draining lymph nodes (tdLN) and nT$^+$ DC are sorted from nTnG steady state draining LN. Cells are co-cultured for 1hr, fixed and analyzed by flow cytometry. (**B**) Quanitfication of coupling (ZsGreen$^+$-nT$^+$ doublets) between ZsGreen$^+$ DC (donor) from tdLN and nT$^+$ DC from steady state LN. Mean coupled frequency is shown. Data are representative of 3 independent experiments.

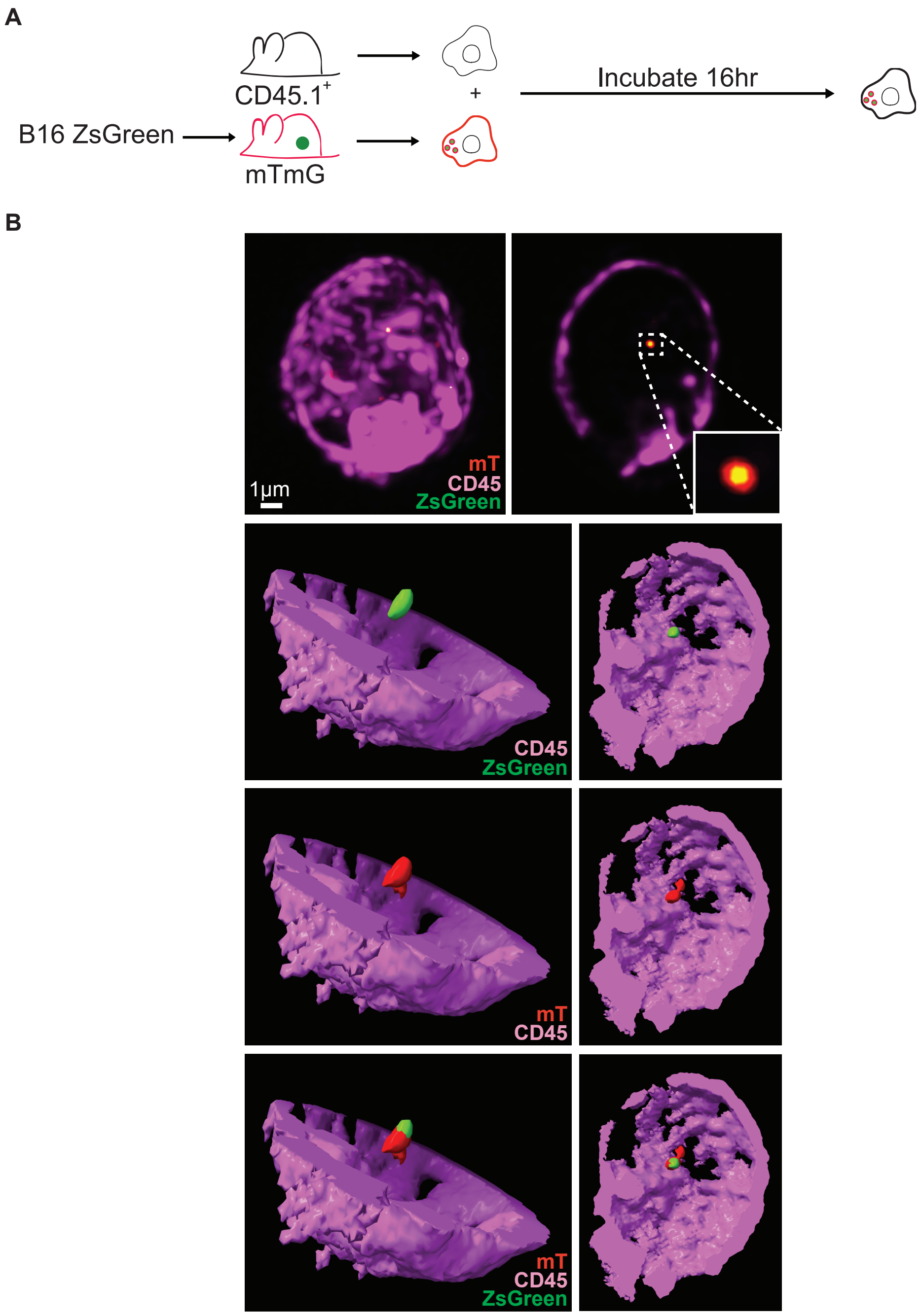


**Fig. S5: Transferred antigens reside in a membrane-bound compartment**
(**A**) Schematic of experimental set up for imaging shown in B. Recipient dendritic cells (DC) and ZsGreen$^{+}$ donor DC are sorted from the lymph nodes (LN) of wildtype, CD45.1$^{+}$ congenic mice and membrane-tdTomato (mT)-expressing mice with subcutaneous B16ZsGreen tumors, respectively. The cells are co-cultured for 16hrs and then ZsGreen$^{+}$CD45.1$^{+}$ DC are sorted and imaged using lattice light sheet (LLS) microscopy. (**B**) LLS imaging of ZsGreen$^{+}$CD45.1$^{+}$ DC obtained from experimental set up shown in A. Vesicle of ZsGreen (green) surrounded by mT (red) derived from mT donor cell membrane within CD45$^{+}$(purple) cell is shown as a maximum intensity projection (MIP) (left, top) and a z-slice (right, top). Surface renderings of the cell show z-slice with additional depth for the cell membrane (CD45) while ZsGreen and mT signal is shown as a surfaced MIP. Representative image.

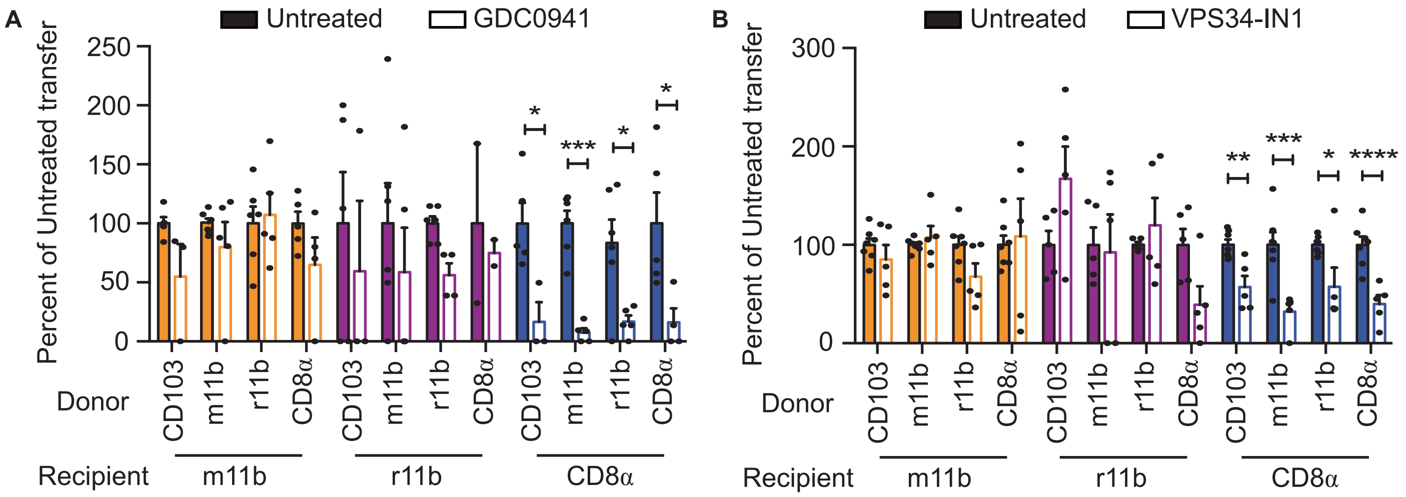


**Fig. S6: Pharmacological inhibition of *in vitro* antigen tranfer**
(**A-B**) Quantification of *in vitro* antigen transfer assay (see Fig. 2C) showing recipient dendritic cell (DC) populations with donor DC in the presence of: (A) GDC0941, (B) VPS34-IN1. Plot shows ZsGreen uptake relative to no inhibitor (untreated, black outlined bars). migratory CD11b DC (m11b), resident CD11b DC (r11b). Error bars represent +/- SEM. Data are representative of 3 independent experiments. *p < 0.05, **p < 0.01, ***p < 0.001, ****p < 0.0001.

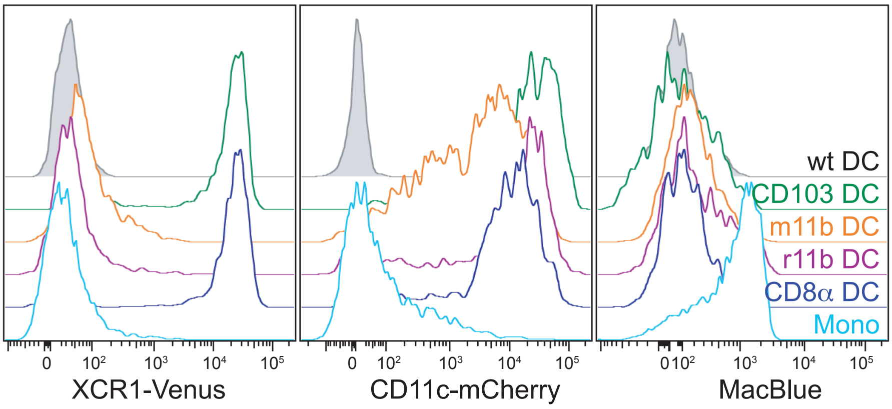


| | CD103 DC | m11b DC | r11b DC | CD8α DC | Mono |
|---|---|---|---|---|---|
| XCR1 | +++ | - | - | +++ | - |
| CD11c | +++ | +++ | +++ | +++ | - |
| MacBlue | - | - | +/- | - | ++ |

**Fig. S7: Reporter expression levels from myeloid cells in the lymph node of XCR1-Venus; CD11c-mCherry; MacBlue reporter mice**
Representative flow cytometric plots showing expression of XCR1-Venus, CD11c-mCherry and MacBlue (CSFR-CFP) within myeloid populations in the tumor-draining lymph node (tdLN). Wildtype (wt), C57BL/6 mouse serves as a control (grey) for reporter expression levels. Data are representative of 2 independent experiments. Summary of gene expression patterns shown in table (bottom). For gating strategy see **Fig. S1**.

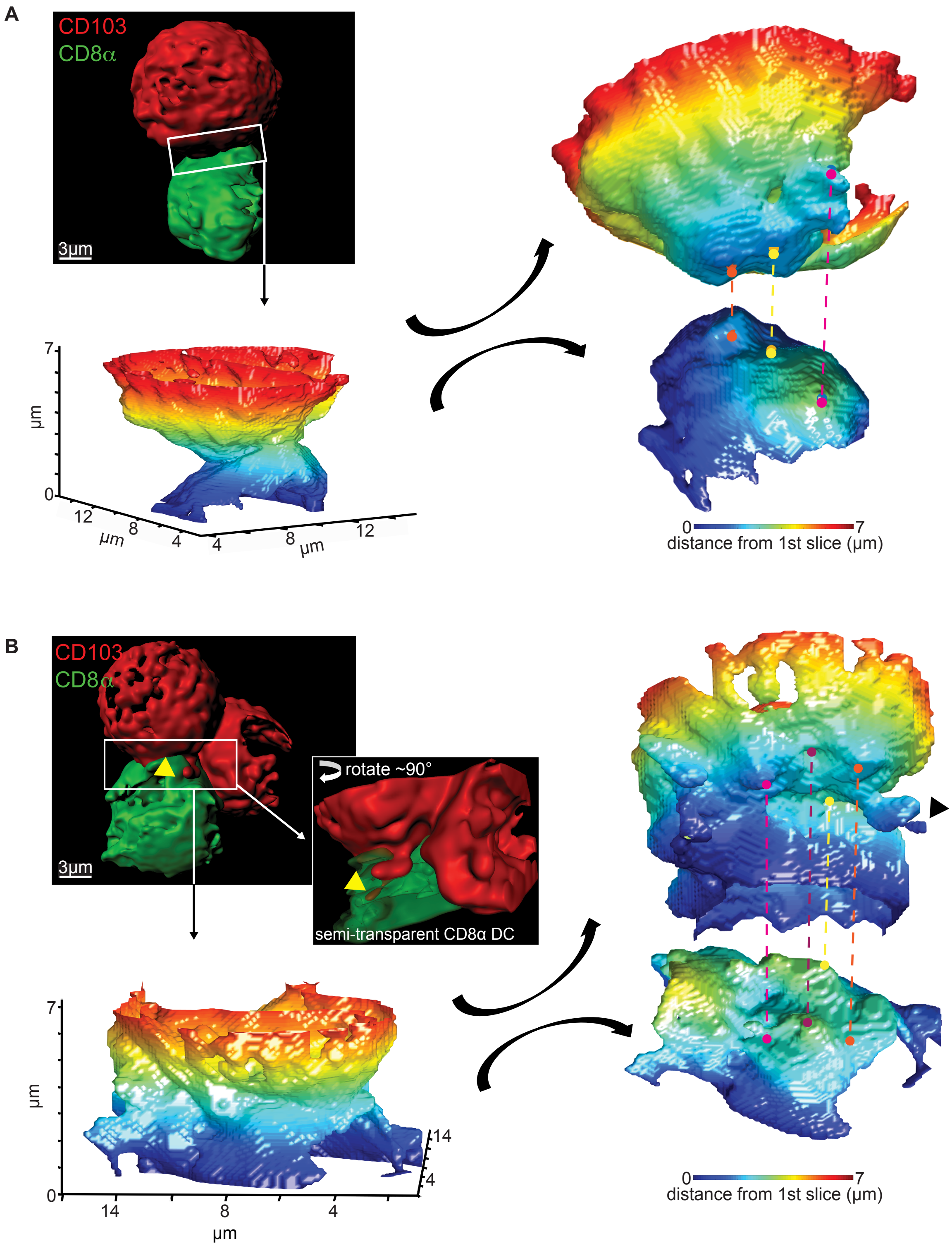


**Fig. S8: Dendritic cell:dendritic cell synaptic characterisation**

(**A**) Example of lattice light sheet (LLS) image of cursory engagement interface of sorted CD103 DC from membrane-tdTomato (mT) mouse and CD8α DC from membrane-GFP (mG) mouse. White box shows interface location used for contact analysis. Heatmap represents distance from reference z-slice (dark blue = 0). Interface is opened (black arrows) and rotated to show surface points of juxtaposed membrane indicated with dots connected with color-coded dotted line. Scale bar = 3μm. (**B**) Example of LLS image of stable, tight engagement interface of sorted CD103 DC from mT mouse and CD8α DC from mG mouse. White box shows interface location used for contact analysis. Yellow arrowheads denote location of finger-like projection of mT membrane into the CD8α DC. Zoomed image shown with semi-transparent CD8α DC surface rotated ~90° to show intercellular mT projection. Heatmap represents distance from reference z-slice (dark blue = 0). Interface is opened (black arrows) and rotated to show surface points of juxtaposed membrane indicated with dots connected with color-coded dotted line. Black arrowhead denotes location of same membrane projection as yellow arrowhead. Scale bar = 3μm.

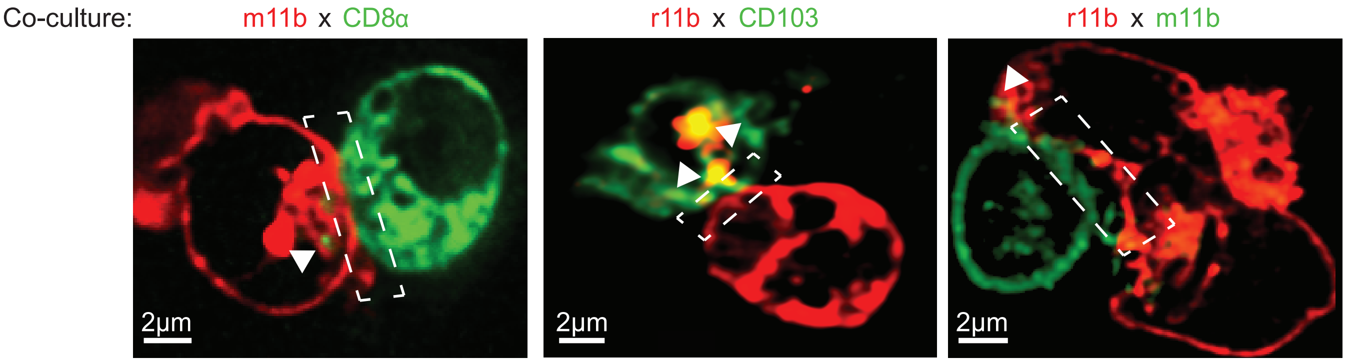


**Fig. S9: Heterotypic interactions occur between various dendritic cell types**
Lattice light sheet imaging of dendritic cells (DC) sorted from lymph nodes (LN) of either membrane-tdTomato (mT, red) expressing or membrane-GFP (mG, green) expressing mice. Images are z-slice depicting interaction interface (white box) between m11b-CD8α (left), r11b-CD103 (middle), or r11b-m11b (right) DC. Arrows indicate membrane exchange event. Images are representative. Scale bar = 2µm.

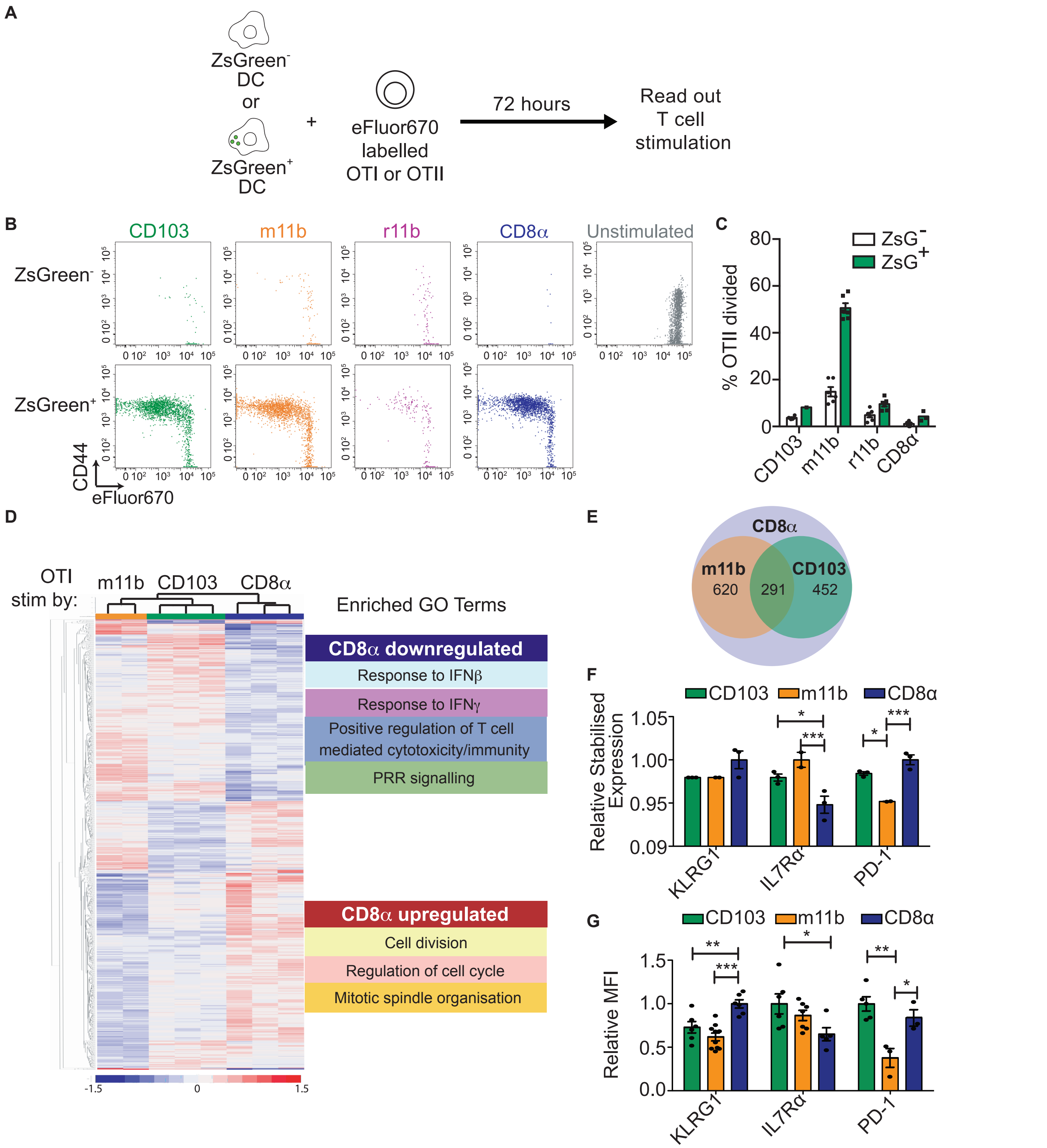


**Fig. S10: T cell stimulation assay using specific dendritic cell subsets**

(**A**) Experimental setup for OTI and OTII, T cell stimulation assay. (**B**) Representative flow cytometric plots showing OTI stimulation as assessed by eFluor670 dye dilution and CD44 upregulation. Example plots are shown for T cells stimulated with each of the 4 different dendritic cell (DC) subsets, with and without ZsGreen, which were sorted from tumor-draining lymph nodes (tdLN) of B16ZsGreen mice. Setup shown in A. (**C**) Percent of CD4$^+$, OTII divided T cells following 72hrs of coculture with different DC types that are either ZsGreen$^-$ (white) or ZsGreen$^+$ (green). Setup shown in A. Plot shows mean % of OTII division based on eFluor670 dye dilution. Error bars represent +/- SEM. Representative of 3 independent experiments. (**D**) Heatmap showing differential gene expression of stimulated CD8$^+$,OTI, T cells detected by RNA-sequencing. Proliferating OTI T cells were sorted following 72hrs of stimulation with ZsGreen$^+$ DC of indicated subtype, orange = migratory CD11b$^+$ DC, green = migratory CD103$^+$ DC, blue = resident CD8α$^+$ DC. GO terms for upregulated and downregulated genes are listed. (**E**) Venn diagram showing differentially expressed genes between T cells stimulated by ZsGreen$^+$ resident CD8α$^+$ DC (purple) compared to migratory ZsGreen$^+$CD103$^+$DC (green) and CD11b$^+$ DC (orange). (**F**) Relative expression of selected genes as determined by RNA sequencing. Error bars represent +/- SEM. Data are 2 experimental replicates for m11b DC and 3 experimental replicates for CD103 and CD8α DC. (**G**) Flow cytometric quantification of protein levels for selected genes shown in D. Plot shows relative mean fluorescence intensity (MFI). Error bars represent +/- SEM. Data are representative of 2 independent experiments. For all graphs $^*p < 0.05$, $^{**}p < 0.001$, $^{***}p < 0.0001$.

| CD8α downregulated | | CD8α upregulated |
|---|---|---|
| Response to IFNβ | Positive regulation of T cell mediated cytotoxicity/immunity | Cell division |
| | | Regulation of cell cycle |
| Response to IFNγ | PRR signalling | Mitotic spindle organisation |

| CD8α Dendritic Cell stimulated GO biological process | Reference List | Genes Downregulated | Expected | Fold Enrichment | P-value |
|---|---|---|---|---|---|
| negative regulation of methylation-dependent chromatin silencing (GO:0090310) | 3 | 2 | 0.02 | > 100 | 4.09E-04 |
| adhesion of symbiont to host (GO:0044406) | 14 | 4 | 0.09 | 43.93 | 4.77E-06 |
| regulation of methylation-dependent chromatin silencing (GO:0090308) | 13 | 3 | 0.08 | 35.48 | 1.39E-04 |
| cytoplasmic pattern recognition receptor signaling pathway (GO:0002753) | 14 | 3 | 0.09 | 32.95 | 1.68E-04 |
| cellular response to interferon-beta (GO:0035458) | 43 | 9 | 0.28 | 32.18 | 4.44E-11 |
| regulation of interferon-gamma-mediated signaling pathway (GO:0060334) | 15 | 3 | 0.1 | 30.75 | 2.01E-04 |
| regulation of response to interferon-gamma (GO:0060330) | 15 | 3 | 0.1 | 30.75 | 2.01E-04 |
| response to protozoan (GO:0001562) | 36 | 7 | 0.23 | 29.9 | 1.08E-08 |
| defense response to protozoan (GO:0042832) | 32 | 6 | 0.21 | 28.83 | 1.53E-07 |
| response to interferon-beta (GO:0035456) | 54 | 10 | 0.35 | 28.48 | 1.04E-11 |
| antigen processing and presentation of endogenous peptide antigen (GO:0002483) | 38 | 7 | 0.25 | 28.33 | 1.50E-08 |
| antigen processing and presentation of endogenous peptide antigen via MHC class I (GO:0019885) | 38 | 7 | 0.25 | 28.33 | 1.50E-08 |
| antigen processing and presentation of endogenous peptide antigen via MHC class I via ER pathway, TAP-independent (GO:0002486) | 33 | 6 | 0.21 | 27.96 | 1.80E-07 |
| antigen processing and presentation of endogenous peptide antigen via MHC class I via ER pathway (GO:0002484) | 34 | 6 | 0.22 | 27.14 | 2.10E-07 |
| antigen processing and presentation of endogenous peptide antigen via MHC class Ib (GO:0002476) | 34 | 6 | 0.22 | 27.14 | 2.10E-07 |
| antigen processing and presentation of peptide antigen via MHC class Ib (GO:0002428) | 35 | 6 | 0.23 | 26.36 | 2.45E-07 |
| antigen processing and presentation of endogenous antigen (GO:0019883) | 41 | 7 | 0.27 | 26.25 | 2.40E-08 |
| antigen processing and presentation via MHC class Ib (GO:0002475) | 39 | 6 | 0.25 | 23.66 | 4.34E-07 |
| antigen processing and presentation of peptide antigen via MHC class I (GO:0002474) | 47 | 7 | 0.31 | 22.9 | 5.58E-08 |
| regulation of interferon-alpha production (GO:0032647) | 28 | 4 | 0.18 | 21.97 | 5.22E-05 |
| regulation of T cell mediated cytotoxicity (GO:0001914) | 55 | 7 | 0.36 | 19.57 | 1.49E-07 |
| innate immune response-activating signal transduction (GO:0002758) | 64 | 8 | 0.42 | 19.22 | 2.11E-08 |
| positive regulation of T cell mediated cytotoxicity (GO:0001916) | 50 | 6 | 0.33 | 18.45 | 1.63E-06 |
| pattern recognition receptor signaling pathway (GO:0002221) | 62 | 7 | 0.4 | 17.36 | 3.14E-07 |
| positive regulation of T cell mediated immunity (GO:0002711) | 71 | 8 | 0.46 | 17.33 | 4.43E-08 |

| CD8α Dendritic Cell stimulated GO biological process | Reference List | Genes Upregulated | Expected | Fold Enrichment | P-value |
|---|---|---|---|---|---|
| mRNA 3'-end processing by stem-loop binding and cleavage (GO:0006398) | 5 | 3 | 0.03 | > 100 | 9.06E-06 |
| attachment of mitotic spindle microtubules to kinetochore (GO:0051315) | 11 | 3 | 0.06 | 49.04 | 5.75E-05 |
| attachment of spindle microtubules to kinetochore (GO:0008608) | 19 | 5 | 0.11 | 47.32 | 1.86E-07 |
| kinetochore organization (GO:0051383) | 17 | 4 | 0.09 | 42.31 | 4.96E-06 |
| histone mRNA metabolic process (GO:0008334) | 17 | 3 | 0.09 | 31.73 | 1.76E-04 |
| microtubule depolymerization (GO:0007019) | 18 | 3 | 0.1 | 29.97 | 2.04E-04 |
| protein localization to chromosome, centromeric region (GO:0071459) | 18 | 3 | 0.1 | 29.97 | 2.04E-04 |
| mitotic metaphase plate congression (GO:0007080) | 41 | 5 | 0.23 | 21.93 | 5.46E-06 |
| metaphase plate congression (GO:0051310) | 52 | 6 | 0.29 | 20.75 | 8.11E-07 |
| mitotic spindle organization (GO:0007052) | 70 | 8 | 0.39 | 20.55 | 1.18E-08 |
| microtubule cytoskeleton organization involved in mitosis (GO:1902850) | 94 | 10 | 0.52 | 19.13 | 3.12E-10 |
| establishment of chromosome localization (GO:0051303) | 70 | 7 | 0.39 | 17.98 | 2.33E-07 |
| chromosome localization (GO:0050000) | 72 | 7 | 0.4 | 17.48 | 2.78E-07 |
| mitotic sister chromatid segregation (GO:0000070) | 98 | 9 | 0.55 | 16.51 | 8.21E-09 |
| regulation of mitotic sister chromatid separation (GO:0010965) | 56 | 5 | 0.31 | 16.05 | 2.22E-05 |
| regulation of chromosome segregation (GO:0051983) | 102 | 9 | 0.57 | 15.87 | 1.14E-08 |
| mitotic nuclear division (GO:0140014) | 134 | 11 | 0.75 | 14.76 | 5.11E-10 |
| regulation of chromosome separation (GO:1905818) | 62 | 5 | 0.34 | 14.5 | 3.50E-05 |
| sister chromatid segregation (GO:0000819) | 126 | 10 | 0.7 | 14.27 | 4.38E-09 |
| mitotic cytokinesis (GO:0000281) | 63 | 5 | 0.35 | 14.27 | 3.76E-05 |
| protein localization to chromosome (GO:0034502) | 63 | 5 | 0.35 | 14.27 | 3.76E-05 |
| aerobic respiration (GO:0009060) | 64 | 5 | 0.36 | 14.05 | 4.04E-05 |
| regulation of mitotic sister chromatid segregation (GO:0033047) | 67 | 5 | 0.37 | 13.42 | 4.96E-05 |
| chromosome segregation (GO:0007059) | 257 | 18 | 1.43 | 12.59 | 1.45E-14 |
| centrosome cycle (GO:0007098) | 75 | 5 | 0.42 | 11.99 | 8.24E-05 |

**Fig. S11: Gene Ontology term enrichment analysis for RNA-sequencing from T cells stimulated by ZsGreen$^{+}$, antigen-bearing, dendritic cells**

Gene Ontology (GO) term enrichment analysis of genes differentially regulated by OTI cells stimulated by resident CD8α$^{+}$ dendritic cells (DC) compared to migratory DC (CD103$^{+}$and CD11b$^{+}$). Analysis performed using common 291 genes shown in overlap of Venn diagram in **Fig. S10E**. Top 25 pathways downregulated (blue) and upregulated (red) in OTI T cells are listed in order of fold enrichment. Color-coded to indicate thematic pathways noted in **Fig. S10D**. Light blue = response to IFNβ, purple = response to IFNγ, dark blue = positive regulation of T cell cytotoxicity, green = pattern recognition receptor signaling, yellow = cell division, pink = regulation of cell cycle, orange = mitotic spindle organisation.

| CD8α downregulated | | CD8α upregulated | |
|---|---|---|---|
| Memory gene sets | Response to IFNγ | Regulation of cell cycle | Early effector gene sets |

| Gene Set Name | Description | p-value | FDR q-value |
|---|---|---|---|
| GSE36527 | Genes up-regulated in KLRG1- SELL low T reg: CD69- versus CD69+. | 2.01E-31 | 2.39E-27 |
| GSE19888 | Genes up-regulated in HMC-1 cells: incubated with ALL1 and treated with Cl-IB-MECA vs stimulation by T cell membranes. | 1.23E-29 | 4.86E-26 |
| HALLMARK_IFNy_response | Genes up-regulated in response to IFNG. | 1.23E-29 | 4.86E-26 |
| REACTOME_Immune_system | Genes involved in Immune System | 7.74E-27 | 2.30E-23 |
| GSE19888 | Genes up-regulated in HMC-1 cells incubated with ALL1 versus those followed by treatment with Cl-IB-MECA. | 3.79E-26 | 9.00E-23 |
| GSE19888 | Genes up-regulated in HMC-1 cells: incubated with ALL1 versus stimulated with T cell membranes. | 1.73E-24 | 3.25E-21 |
| GSE33424 | Genes up-regulated in CD8 T cells: KLRB1 int versus KLRB1-. | 1.91E-24 | 3.25E-21 |
| GSE13485 | Genes down-regulated in unstimulated PBMC versus PBMC 7 days after stimulation with YF17D vaccine. | 9.04E-23 | 1.07E-19 |
| GSE26890 | Genes up-regulated in effector CD8 T cells: CXCR1+ versus CXCR1-. | 9.04E-23 | 1.07E-19 |
| GSE43863 | Genes up-regulated in CD4 SMARTA effector T cells during acute infection of LCMV: follicular helper (Tfh) versus Ly6c low CXCR5-. | 9.04E-23 | 1.07E-19 |

| Gene Set Name | Description | p-value | FDR q-value |
|---|---|---|---|
| GOBERT_OLIGODENDROCYTE_DIFFERENTIATION_UP | Genes up-regulated during later stage of differentiation of Oli-Neu cells in response to PD174265. | 4.50E-57 | 5.36E-53 |
| FISCHER_DREAM_TARGETS | Target genes of the DREAM complex. | 5.22E-49 | 3.10E-45 |
| GSE13547 | Genes up-regulated in B lymphocytes: control versus stimulated by anti-IgM for 12h. | 2.14E-43 | 8.48E-40 |
| FLORIO_NEOCORTEX_BASAL_RADIAL_GLIA_DN | Genes down-regulated in bRG relative to aRG, and up-regulated in both aRG and bRG relative to neurons. | 7.76E-43 | 2.31E-39 |
| MARSON_BOUND_BY_E2F4_UNSTIMULATED | Genes with promoters bound by E2F4 in unstimulated hybridoma cells. | 7.35E-41 | 1.75E-37 |
| GOLDRATH_EFF_VS_MEMORY_CD8_TCELL_UP | Genes up-regulated in comparison of effector CD8 T cells versus memory CD8 T cells. | 3.31E-40 | 6.56E-37 |
| KINSEY_TARGETS_OF_EWSR1_FLII_FUSION_UP | Genes up-regulated in TC71 and EWS502 cells by EWSR1-FLI1 as inferred from RNAi knockdown of this fusion protein. | 1.83E-38 | 3.10E-35 |
| GSE15750 | Genes up-regulated in comparison of wild type CD8 effector T cells at day 6 versus those at day 10. | 2.90E-38 | 4.31E-35 |
| MODULE_54 | Cell cycle (expression cluster). | 7.64E-37 | 1.01E-33 |
| GSE15750 | Genes up-regulated in comparison of wild type CD8 effector T cells at day 6 versus those from mice deficient for TRAF6 at day 10. | 2.40E-36 | 2.60E-33 |

**Fig. S12: Molecular signature database enrichment analysis for RNA-sequencing from T cells stimulated by ZsGreen$^+$, antigen-bearing, dendritic cells**
Molecular signature database enrichment analysis of genes differentially regulated by OTI cells stimulated by resident CD8α$^+$ dendritic cells (DC) compared to migratory DC (CD103$^+$and CD11b$^+$). Analysis performed using common 291 genes shown in overlap of Venn diagram in **Fig. S10E**. Pathways downregulated (blue) and upregulated (red) in OTI T cells are listed in order of fold enrichment. Color-coded to indicate thematic pathways. Light blue = gene sets involved in memory T cell differentiation, purple = response to IFNγ, pink = regulation of cell cycle, orange = gene sets involved in early effector T cell differentiation.
